# Cell-Driven Fluid Dynamics: A Physical Model of Active Systemic Circulation

**DOI:** 10.1101/2024.05.19.594862

**Authors:** Yufei Wu, Morgan A. Benson, Sean X. Sun

## Abstract

Active fluid circulation and transport are key functions of living organisms, which drive efficient delivery of oxygen and nutrients to various physiological compartments. Because fluid circulation occurs in a network, the systemic flux and pressure are not simple outcomes of any given component. Rather, they are emergent properties of network elements and network topology. Moreover, consistent pressure and osmolarity gradients across compartments such as the kidney, interstitium, and vessels are known. How these gradients and network properties are established and maintained is an unanswered question in systems physiology. Previous studies have shown that epithelial cells are fluid pumps that actively generate pressure and osmolarity gradients. Polarization and activity of ion exchangers that drive fluid flux in epithelial cells are affected by pressure and osmolarity gradients. Therefore, there is an unexplored coupling between the pressure and osmolarity in the circulating network. Here we develop a mathematical theory that integrates the influence of pressure and osmolarity on solute transport and explores both cell fluid transport and systemic circulation. This model naturally generates pressure and osmolarity gradients across physiological compartments, and demonstrates how systemic transport properties can depend on cell properties, and how the cell state can depend on systemic properties. When epithelial and endothelial pumps are considered together, we predict how pressures at various points in the network depend on the overall osmolarity of the system. The model can be improved by including physiological geometries and expanding solute species, and highlights the interplay of fluid properties with cell function in living organisms.

---

Fluid circulation across various physiological compartments is driven by overall blood flow and active pumping of solutes across epithelial and endothelial barriers. The overall circulation in humans in large (100s of liters per day), and can be considered essentially as a closed circuit. For example, in the kidney, fluid transport from the lumen side (apical) to the interstitial side (basal) is driven by passage of mostly Na^+^ and Cl*^−^* across the epithelial layer by ion transporters [1, 2] (Fig. 1). Because water follows solute transport, small concentration gradients of solutes will drive water flow and generate hydraulic pressure gradients. On the other hand, it was shown that the presence of hydraulic pressure gradients can influence cell ion channel apical-basal polarization and change solute/water flux [3]. Thus, pressure gradients can directly influence solute flux at the cellular level. Experimental data on the pressure dependence of solute and water transport has been available in the literature for epithelial cells [4, 5, 6, 7], endothelial cells [8] and possibly others.

**Figure 1.**
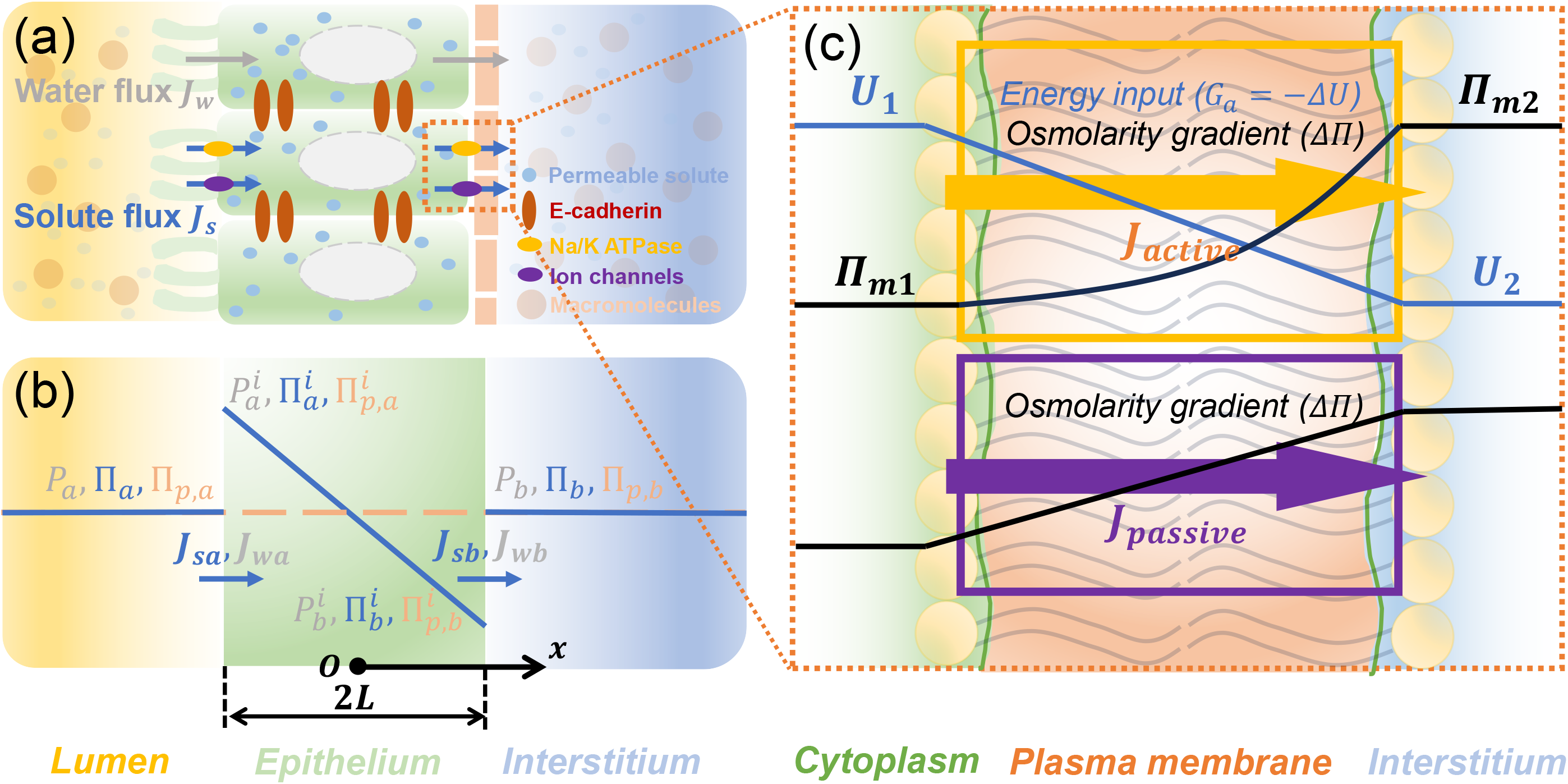
(a) An illustration of active water transport in epithelial layer. Water transport is driven by gradients of hydraulic pressure and osmotic pressure across the apical and basal cell membrane. The osmotic pressure arises from impermeable macromolecules (e.g., proteins) and permeable molecules (e.g., NaCl). The osmotic pressure gradient of permeable solute is established by both active ion pumping (e.g., Na/K ATPase) and passive ion transport (ion channels). (b) A diagram of the model (details in SM) and predicted spatial distribution of hydraulic pressure (*P*), osmotic pressure of permeable molecules (Π) and impermeable macromolecules (Π*_p_*). Subscripts *a, b* denote external apical and basal side of the epithelium,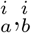 denote apical and basal side of the cell interior. *J_w_* and *J_s_* denote water and solute flux, respectively. (c) Permeable solute flux across the membrane results from active pumping and passive diffusion. Active pumping is driven by a free energy function *U* across the cell membrane. Passive solute flux is driven by the solute concentration gradient. Π*_m_* is the predicted solute concentration profile within the membrane (SM).

Pressure natriuresis is also a well-known phenomena driven by blood pressure-dependence of water transport in the kidney [9, 10]. The complex interplay between solute concentration, oncotic pressure, interstitial and blood hydraulic pressures in systemic circulation have not been considered theoretically. Moreover, pressure and osmolarity differences in the interstitium, vessels and capillaries, the lymphatic space, and subcutaneous compartments are known [11, 12, 13, 14], which must be emergent properties of the overall circulation network. These facts suggest that a unified mathematical framework that starts with cell properties and ends with systemic fluid circulation predictions is needed. In this paper, we develop a mathematical model of systemic fluid circulation, incorporating active pumping properties of epithelial and endothelial cells and network connectivity. The model starts with cell transport properties, and incorporates pressure- and osmolarity-dependent solute transport at the single cell level. We then use the cell scale model to derive transport/pumping properties of epithelial and endothelial barriers. The results directly relate phenomenological coefficients of transport equations with cell-level properties. We then incorporate active pumping properties of cells in a circulation network model, but now include both solute and fluid pumping in the network. The model predicts overall solute and fluid flux in the network as well as osmolarities and pressures in the various compartments in the network. Gradients of osmolarity and pressure are natural outcomes of the model, and can influence each other. We show that inclusion of active transport or pumping properties of endothelia/epithelia fundamentally changes the overall network circulation properties. Specifically, the model predicts how the total osmolarities of the system can influence network transport properties and pressure/osmolarity gradients across compartments. We also point out how the model can be extended to include realistic network geometries and physiology-level feedback control. Mechanical rigidity of the network can be relaxed and growth can be included to allow the network morphology to adapt to pressure/osmolarity changes, leading to a fluid-centric theory of morphogenesis.

## Cell monolayer fluid and solute pumping model

The first element is to develop a model of a single epithelial/endothelial layer where fluid and solute transport are treated together. Details of this model are given in the supplemental material (SM). The model considers a single neutral solute (e.g., NaCl) driven by active solute transporters. This is a simplification that does not include complexities of multiple ionic species and their possible interaction with electrical fields. A more detailed model with chemical complexity can be included in a followup model. In addition to small permeable solutes, we also include impermeable macromolecules such as proteins, which do not move through the membrane. The osmotic pressure of these macromolecules is denoted by Π*_p_*. The cell layer is modeled as a domain of thickness 2*L* (Fig. 1), where solute and water fluxes, *J_s_* and *J_w_*, can occur at the two surfaces facing the lumen and the interstitium, which have osmotic pressures and hydraulic pressures (Π*_a_,* Π*_b_,* Π*_p,a_,* Π*_p,b_, P_a_, P_b_*), respectively. The primary principle governing these flows is the continuity of fluxes. In other words, *J_s_* and *J_w_* across the surfaces are continuous with respect to the fluxes in the cytoplasmic domain. The permeable solute flux in the cytoplasmic domain is a combination of diffusion plus convection:

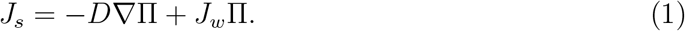

where Π = *RTc* is the osmotic pressure, *c* is the solute concentration. *D* is the solute diffusion coefficient and *RT* is the gas constant times temperature. For convenience, we incorporate the factor of *RT* into solute flux *J_s_*. The water flux in the cytoplasm, *J_w_*, contributes to the convection of the solute. It is related to the hydraulic pressure gradient in the cytoplasm: *J_w_ ∝ ∇P* (SM). Therefore the solute and water flux must be solved together in a coupled manner. In 1D, water flux (or velocity) in the cytoplasm becomes: 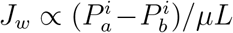. Here, *μ* is the dynamic viscosity of fluid. 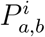 is the hydraulic pressure just inside the cell at the apical/basal surface.

To describe the ’pumping’ behavior of solutes driven by active ion transporters and passive permeation, a model of permeable solute flux across the cell membrane is needed (Fig. 1c). In general, near equilibrium, flux of material is proportional to the free energy gradient and material concentration *c*: *J_s_ ∝ c∇F*, and the free energy function across the membrane is *F* = *RT* ln *c* + *U*, where *U* models an energy input by the cell. Solute is transported via two different types of channels: For passive ion channels, *U* = 0 and there is no energy input. For active ion pumps, *U* ≠ 0, and the energy difference across the membrane, *G_a_* = *−*Δ*U* = (*U*_2_ *− U*_1_) in Fig. 1c, is the driving force of the active solute transport. Thus, in our model, a single parameter, *G_a_*, describes the active solute transport properties of the cell.

Inside the membrane domain, the solute flux is continuous. Adding the fluxes through both types of solute carriers together, we obtain the total flux: 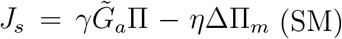. Here 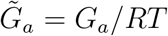 and ΔΠ*_m_* is the osmolarity difference across the membrane. *γ* is the active ion transport coefficient, and *η* is defined as the sum of active and passive ion transport coefficients (SM). Similarly, the water flux across the cell membrane is proportional to water free energy difference across the membrane: *J_w_* = *α*(Δ*P −* ΔΠ *−* ΔΠ*_p_*) [15]. Note that the macromolecules (Π*_p_*) contribute to water flux *J_w_* but do not directly contribute to the solute flux *J_s_*. Also, the solute flux depends on the total osmolarity Π, not just the osmolarity difference ΔΠ*_m_*. This is a direct result of the free energy function across the membrane, and will have important implications for systemic flux and pressure later. In the following, when not specified, the solute osmotic pressure (Π) only refers to that of the permeable small molecules.

The parameter *G_a_* describes the active solute transport property of the cell. Experiments have shown *G_a_* is not a constant, but can depend on pressure. When pressure is applied to the basal side of a kidney epithelium, sodium/potassium exchanger (NKE) is seen to leave the basal-lateral side, slowing the overall solute flux [3]. NaCl flux across the epithelium is the major driver of water flux and NaCl flux also has been shown to decline with increasing pressure [5]. Therefore, cells reduce the solute driving force when pressure at the basal side is increased. Similar arguments can be applied to osmotic pressure, although direct experiments are still needed. A simple model that incorporates this pressure/osmolarity sensing by cells is: 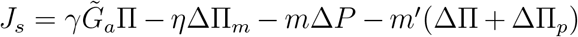, where (*a, b*) denote apical and basal surface, Δ*P* = *P_b_ − P_a_,* ΔΠ = Π*_b_ −* Π*_a_,* ΔΠ*_p_* = Π*_p,b_ −* Π*_p,a_* are the differences across the cell. Here we assume that the cells are sensing the total osmotic pressure contributed by both small solutes and impermeable macromolecules. *m* and *m′* describe solute transporter polarization as a function of osmolarity and pressure gradients across the membrane.

Once the water and solute fluxes across the apical and basal membranes are determined, the fluid and solute transport equations can be solved across the epithelium simultaneously to obtain the overall concentration profile and the pressure field (SM). An excellent analytic approximation can be made (SM) and the results are

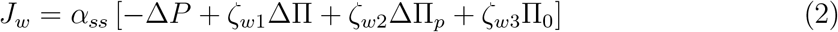

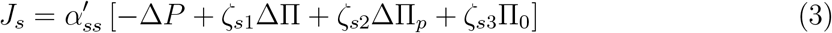

where *J_w_* (*J_s_*) is the water (solute) flux across the epithelium, respectively. ΔΠ = Π*_a_ −*Π*_b_*, ΔΠ*_p_* = Π*_p,b_−*Π*_p,a_*, Δ*P* = *P_a_−P_b_* and 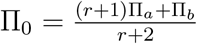 is a weighted mean osmotic pressure of the permeable molecules. 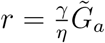 is the dimensionless energy input by the cell, which is the relative capacity of active ion transport compared to total ion transport. Note *r* ≪ 1, and therefore Π_0_ *∼* (Π*_a_* + Π*_b_*)*/*2. 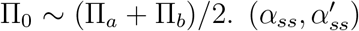 are the two effective permeability coefficients which are functions of molecular parameters. *ζ_wi_, ζ_si_*(*i* = 1, 2, 3) are also functions of cell/molecular parameters. In particular, *ζ_si_* also depends on the mean osmotic pressure of permeable molecules (Π_0_) (SM). Eqs. (2,3) represent the so called pump performance curve (PPC), which describes how flux changes with external pressure. We can also include the paracellular (through junctions) water and solute fluxes, which only modify the coefficients in Eqs. (2, 3). Detailed discussion, including a glossary of parameters, are given in the SM.

The computed solute osmolarity profiles in the cytoplasm are shown in Fig. 2(a)-(b), which demonstrate how the cytoplasmic concentration profiles can adapt to the external osmolarity for a given energy input and pressure gradient, captured by the dimensionless cell energy input 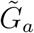 and Δ*P*. As the external osmolarity and pressure gradients change, the cytoplasmic osmolarity profiles also vary following the external change. There is a critical osmolarity (pressure) difference such that the flux can be reversed. The energy input, and other cell properties such as membrane permeability can be adjusted by the cell by changing polarization/gene expression. Eq. 2 is consistent with the classic Starling’s equation of endothelial leakiness: *J_w_* = *α*(*−*Δ*P* + *σ*ΔΠ) [16]. In the limit 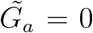 and no osmolarity difference for permeable solutes (ΔΠ = 0, ΔΠ*_p_ /*= 0), the predicted Starling coefficient is 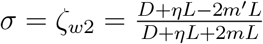.

**Figure 2.**
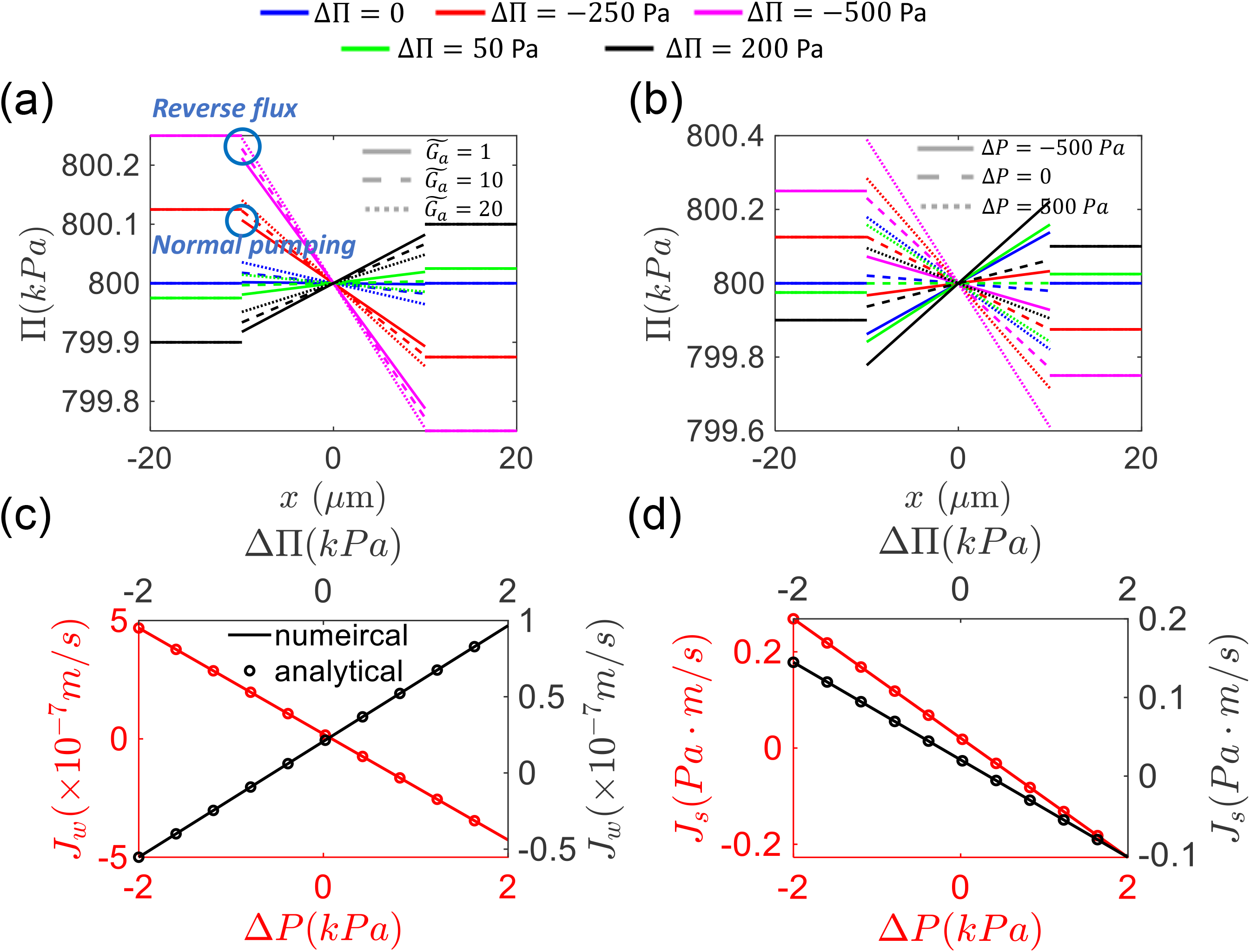
(a)-(b) Cytoplasmic solute concentration (osmolarity) profile when external osmolarity and pressures conditions are varied. The osmolarity profile in the cytoplasm adapts to external changes. Water flux decreases when external basal-apical osmotic pressure difference increases, and the transport direction could reverse. Increasing the energy input and the pressure gradient decrease the slope of the inner concentration profile. Here, the mean osmotic pressure of basal and apical sides are 800 *k*Pa, and only ΔΠ is changed. In (a) and (b), *m* = *m^t^* = 0. (c)-(d) Comparisons between the accurate numerical solution and the analytic approximation for water and solute flux. In the calculation, when not specified, Δ*P* and ΔΠ are set as zero and the mean osmotic pressure is set as Π_0_ = 800 *k*Pa.

The results for solute and water fluxes predict that cells can adapt to changing osmolarity and pressure conditions (Fig. 2(c)-(d)). These fluxes are consistent with the idea of active ”pumping”, i.e., the flux declines with increasing hydraulic pressure difference: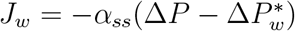 and 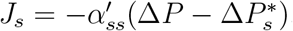, where 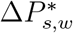 are the stall pressures when fluxes reach 0. When the external gradients are too high, the flux can be reversed.

Interestingly, we predict that the total background osmolarity, Π_0_, has an effect on water and solute fluxes. Depending on the pressure and osmolarity gradient, the mean external osmolarity may have opposite influence on the fluxes (Fig. 3(a)-(b)). The critical condition is derived in the SM. With a sufficiently high osmolarity, water flux is limited by the rate of active ion pumping (*γ*) and energy input 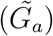. The limiting water flux when mean osmolarity reaches infinity is predicted to be 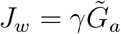 by Eq. S24.

**Figure 3.**
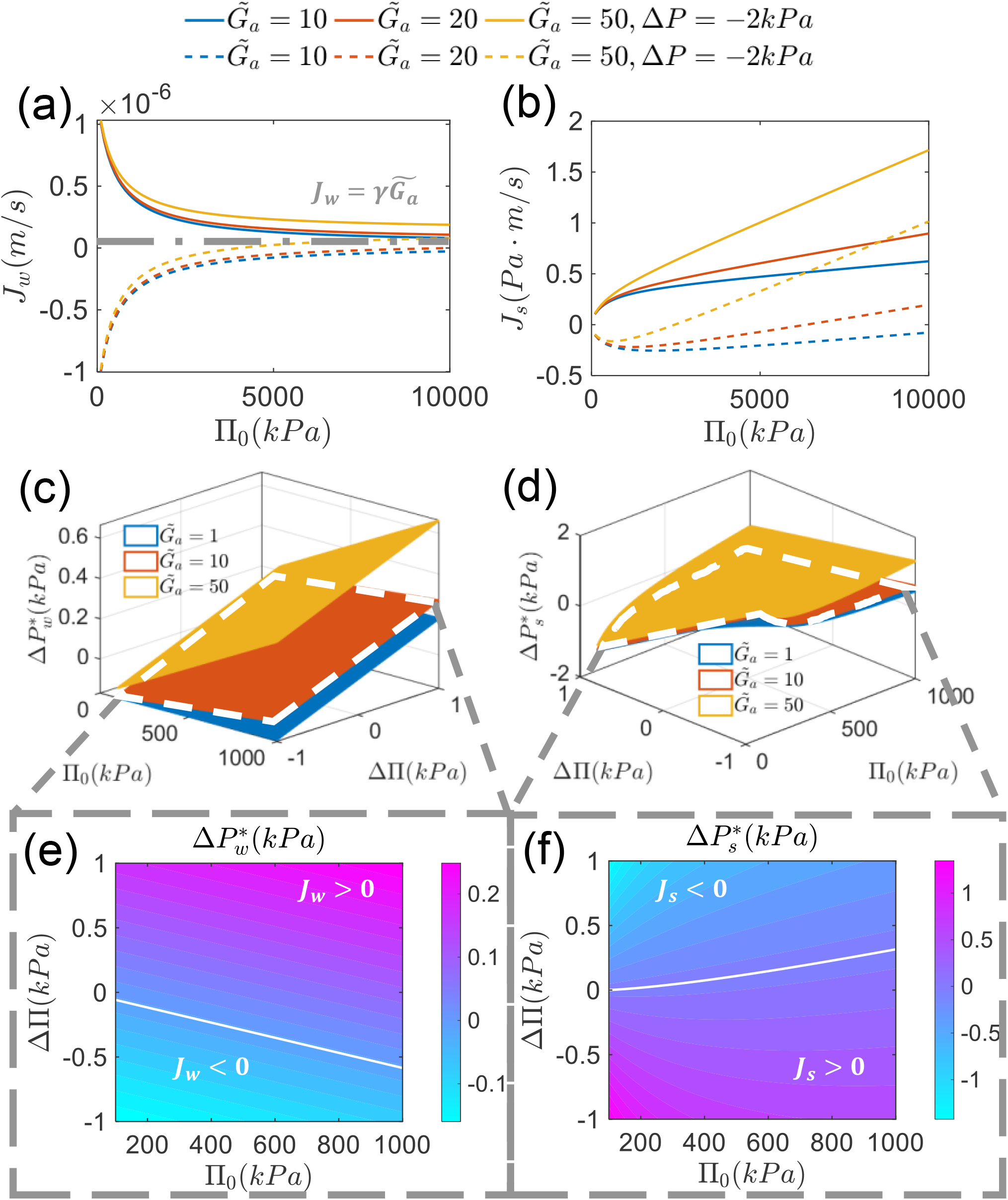
(a)-(b) Influence of mean background osmotic pressure Π_0_ on water and solute flux. There are two regimes in the phase diagram. The boundary is determined by the pressure gradient (Δ*P*), background osmotic pressure (Π_0_) and osmotic pressure gradient (ΔΠ, ΔΠ*_p_*). There is a limit for water flux 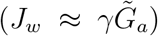 when the osmolarity Π_0_ is sufficiently high. (c)-(d) Dependence of stall pressure for both water 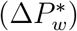 and solute 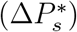 on absolute osmolarity (Π_0_), osmolarity gradient (ΔΠ) and energy input 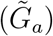. (e)-(f) Phase diagram of water and solute transport. The boundary is the stall pressure, across which the transport direction reverses. For the water flux, the pump functions normally above the boundary line and “reverses” below the boundary line. For solute flux, the region below is the normal regime. As the basal-apical pressure difference (Δ*P*) increases, reversed flux becomes more likely for both water and solute.

Our model can also predict stall pressure/osmolarity (zero flux) as well as phase diagrams of when flux reversals can occur (Fig. 3(c)-(f)). The stall pressures are:

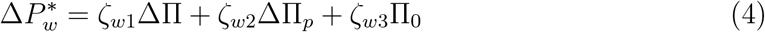

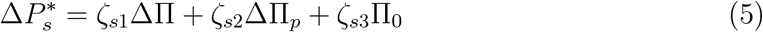

The stall pressures depend on external osmolarity and ion pump relocalization. Pressure or osmolarity changes will result in cell polarization change and ion pump re-localization. This is described by coefficients *m* and *m′* (included in coefficients *ζ_wi_, ζ_si_*). Note that epithelial mechanical integrity may also ultimately determine stall pressure, but this is not captured here. In the results above, ΔΠ refers to permeable solute osmolarity difference across the pump. The osmolarity difference of impermeable macromolecules is set as zero (ΔΠ*_p_* = 0).

## Connected Microphysiological Organ Systems: A Model of Fluid Circulation

In the previous section we discussed an isolated pumping unit. In an organism, epithelia and endothelia are components in a complex circulatory system. The overall fluid circulation flux is a systemic property with central importance in physiology and medicine. Therefore it is of great interest to examine the effect of ”pumping” on the circulatory network from a systemic viewpoint. Modern bioengineering are developing microphysiological systems, where microfluidics are combined with organoid culture to study biological transport and function [17, 18, 19]. These systems allow for microscale control of pressure and osmolarity, which can reveal cell-driven transport properties. There are also many simulation studies on the human circulatory system using lumped parameter models [20, 21, 22, 23]. In analogy to electrical circuits, models such as the Windkessel model treat fluxes in blood vessels using concepts of resistors and capacitors [24]. Despite detailed modelling of pressures in the network, the influence of osmotic pressure and solutes is generally neglected. Moreover, no ”pumping” elements are included, which significantly alters the local and global properties of the circulatory system. Here, we will use the lumped parameter approach, but develop a physiological model that includes possible epithelial/endothelial pumping units. We seek to understand local and global influence of the pump on the overall network. We also explore the influence of osmolarity and energy input of the pump on the overall circulation.

The model consists of three elements: the blood vessel (mainly aorta and vena cava), capillary blood vessels in the organs (both resistance and compliance are considered) and the pumping element (epithelial and endothelial tissues). The pumping element is modeled using water and solute fluxes given in Eqs. (2-3). All blood vessels (aorta, vena cava and organ capillaries) are described by a two-element Windkessel model, which is a resistor-capacitor set in parallel.

## A one-pump circulation model

We first examine a one-pump circuit model in which the pumping element incorporates the epithelial layer, endothelial layer, and interstitium. The inlet of the pump is lumen side of the renal tubule and the outlet faces the lumen of blood vessels. An illustration of the model is shown in Fig. 4(a). Blood is pumped from the heart (modeled as a source with pressure *P_s_*) and goes through aorta (with resistance *R_A_*), and branches before the kidney. A portion of this flow goes through the capillary blood vessel of the organs (effectively modeled as a resistor *R_O_*) while others are filtered by the glomerulus (*R_G_*) and then goes through the pumping unit consisting of epithelium, interstitium and endothelium (water reabsorption). The two branches finally merge and return to the heart through the vena cava (*R_V_*). For convenience, we define the node pressure at the end of vena cava to be zero. All the capacitors can be combined together as one effective capacitor *C_E_*, which gives trivial prediction on dynamical response of the pressure and flux. Therefore, we will neglect ”capacitors” and focus on the static property of the system.

**Figure 4.**
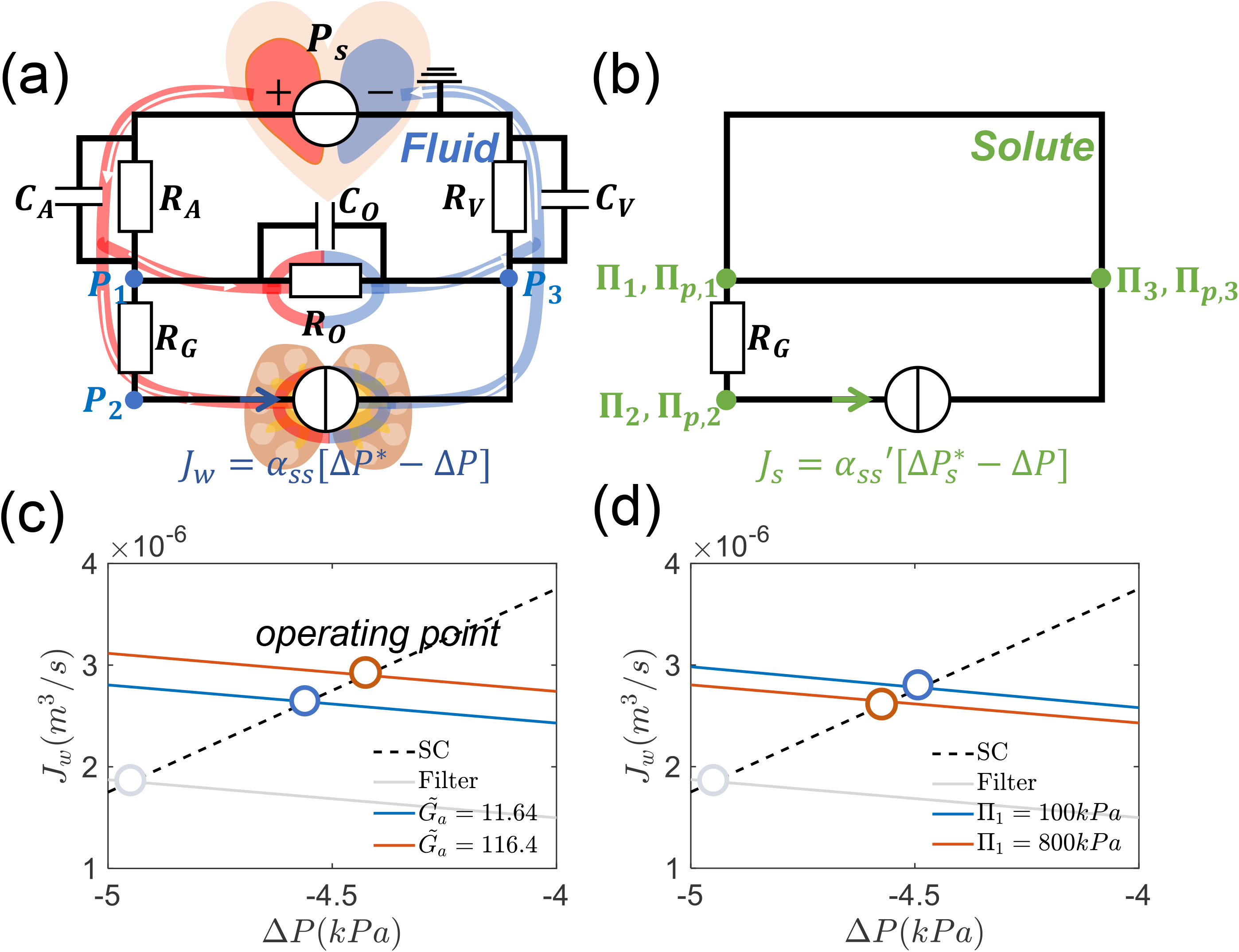
(a) Diagram of the one-pump circulatory system. The circulatory system is simplified into 6 parts: heart aorta, vena cava, organs, glomerulus and the pumping element including renal tubule epithelial tissue and endothelial tissues. Each part is modeled as a two-element Windkessel element. (b) The solute circuit is considered simultaneously with the fluid circuit. The total osmotic pressure (Π+Π_*p*_) is different across the glomerulus *R*_*G*_ (Π_1_ = Π_3_ ≠ Π_2_). (c)-(d) The systems curve, pump performance curve and the operating point. Increasing the cell energy input elevates flux and pressure across the pump while the external osmotic pressure decrease flux and pressure. When the pumping element is replaced by a passive filter 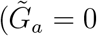, both flux and pressure gradient decrease significantly. In (c), the oncotic pressure (Π_*p*_) in the blood plasma is fixed and the osmotic pressures at different nodes always satisfy: Π_1_ = Π_2_ = Π_3_, Π_*p*,1_ = Π_*p*,3_, Π_*p*,1_ − Π_*p*,2_ = Π_*p*_.

Similarly, we can define the circuit of osmolytes, in which the hydraulic pressure is replaced by osmotic pressure (Fig. 4(b)). The osmotic pressure remains the same through the blood capillaries in organs but changes across the glomerulus. The plasma proteins are not filtered through the glomerulus and therefore create an oncotic pressure difference Π_*P*_, which is approximately Π_*p*,1_ − Π_*p*,2_ = Π_*P*_ = 3.8*kPa* [25]. In the one-pump model, the osmotic pressures of permeable solutes are equal at all nodes.

Just as in an electrical circuit, we can use Kirchhoff’s law to compute pressures at different nodes. However, different from Ohm’s law, the water flux through an element (e.g., capillaries in the organ, pumping element) is a function of both hydraulic pressure and osmotic pressure differences across the element, which is: *J_w_* = (Δ*P −* ΔΠ *−* ΔΠ*_p_*)*/R*, where *R* is the “effective resistance” of the element. Osmolarity is constant in the vessels and only changes across the glomerulus and the pumping element. Consequently, the solute flux in vessels is only proportional to the water flux. The solute flux through a pumping element is given by Eq. 3. There are three nodes in the network and the equations are:

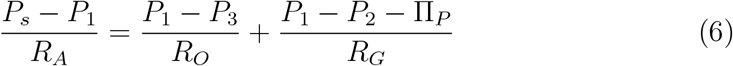

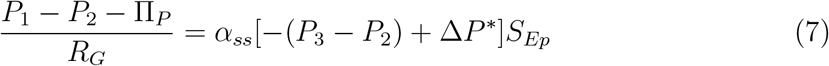

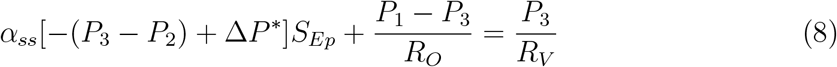

where *R_A_, R_O_, R_G_, R_V_* are resistances of aorta, capillaries in organs, glomerulus and vena cava, respectively. *P_s_* is the pressure heart generates and *P*_1_*, P*_2_*, P*_3_ are the node pressures. Π_1_, Π_3_ are osmotic pressure at nodes 1 and 3. The pumping performance of the kidney cells is: *I* = *α_ss_*[Δ*P ^∗^ −* (*P*_3_ *− P*_2_)], where *α* is a permeability constant and Δ*P ^∗^* is the ”stall pressure” given by eq. 4. The coefficient *S_Ep_* is the total surface area of the renal tubule, which transforms the velocity (m/s) into the volume flow rate (m^3^/s). By defining *R_O_* = *R, R_A_* = *k*_1_*R, R_G_* = *k*_2_*R, R_V_* = *k*_3_*R*, we can get general analytical solutions for all the node pressures, total flux and branch flux (SM).

An important aspect of the circuit is the systems curve, which determines the flux and pressure drop across the pump. The intersection of the pump performance curve and the systems curve is the operating parameters of the pump when it is placed into the circuit. The systems curve is derived from Eqs. 6 - 8:

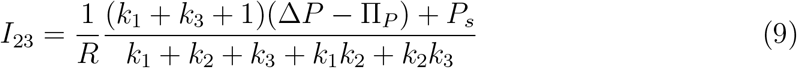

Figure 4 (c)-(d) show the systems curve and the PPC with different energy input 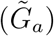 and the osmolarity of the permeable solute (Π_1_). With increased energy input, the pump performance curve shifts upward, resulting in increased water flux. With increase Π_1_, however, the systemic flux decreases. When active pumping is removed 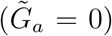, both the flux and pressure difference decrease significantly. The results are intuitive, but we see that systemic properties such as resistances in various vessels will impact flow and pressure across the epithelium.

Fig. S5 systematically explores how the energy input 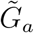, blood oncotic pressure (Π*_p_*) and external osmotic pressure (Π_0_) of permeable solutes influence node pressures and water flux (SM). In particular, the oncotic pressure results provide some explanations of kidney diseases related to protein filtration defects. If the glomerulus cannot block blood proteins, osmotic pressure is increased at node 2 and changes water flux and pressure distribution.

## A two-pump circuit model including the interstitium

In the previous section, epithelial tissue and endothelial tissue are combined together with interstitium to form an equivalent pump. In reality, there are osmolarity and pressure differences across epithelial/interstitial/endothelial compartments, generated by active solute transport [26, 27]. Differences in osmotic pressure and hydraulic pressure in three compartments potentially play an important role in water transport. Additionally, although data is inconclusive, endothelial cells could also actively transport ions and water [28, 29]. In this section, we explore a more detailed model which includes the epithelial pump, possible endothelial pump, and the interstitium. Fig. 5 (a)-(b) show the circuit in consideration. Node 3 corresponds to the interstitium. We solve the coupled system that includes both the hydraulic pressure circuit and osmotic pressure circuit. Using Kirchoff’s law, pressures at four nodes are:

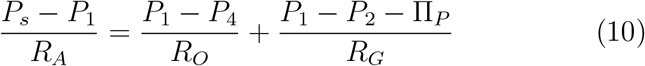

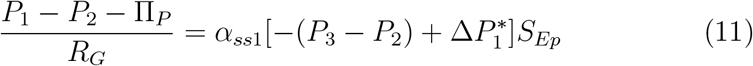

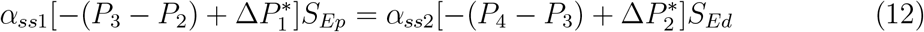

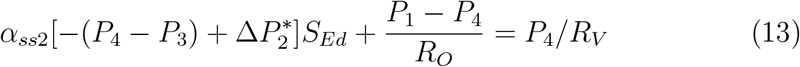

where 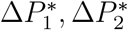 are stall pressures for epithelial and endothelial cells, respectively. To obtain a solution, another equation describing solute flux balance in the interstitium is needed (*J*_*s*1_ = *J*_*s*2_), which can be obtained from Eq. 3.

**Figure 5:**
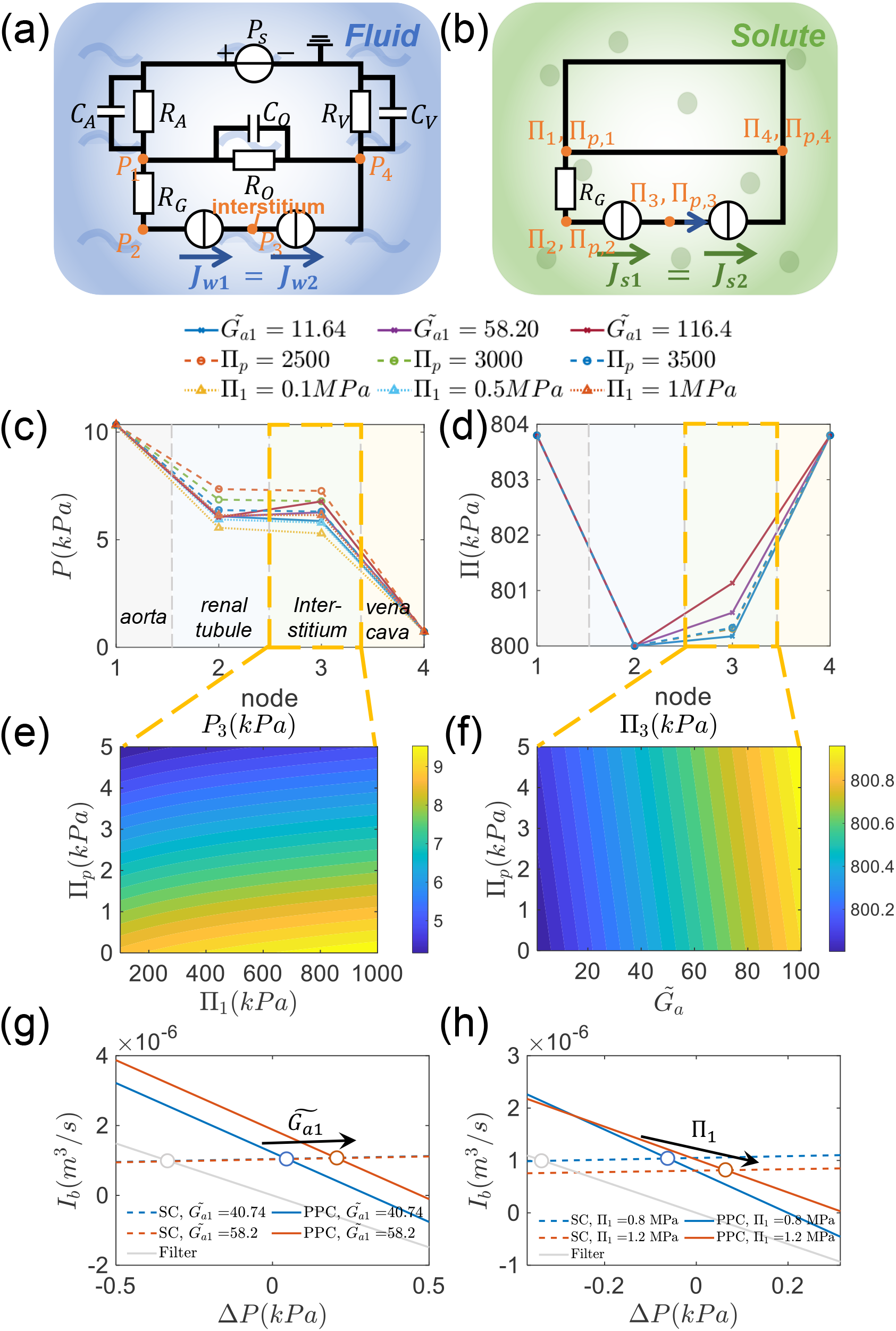
(a)-(b) Diagram of the water and solute circuit for the two-pump model. (a) Fluid circuit. Different from the one-pump model, the epithelial tissue, endothelial tissue and interstitium are all included. (b) Solute (osmolarity) circuit. The osmotic pressures are the same across the organs (node 1 and 4). There is an osmolarity gradient after the glomerulus since proteins filtered. The osmotic pressure difference includes oncotic pressure. Osmotic pressure at node 3 may also be different, which is determined by the solute flux balance. (c)-(d) spatial distribution of pressure and osmolarity. (e)-(f) Influence of total blood osmotic pressure, oncotic pressure and energy input on hydraulic pressure and osmotic pressure in the interstitium.(g)-(h) The systems curve, the pump performance curve, and the operating point corresponding to the ”epithelial pump”. In all calculations, when not specified, the energy inputs for kidney epithelial pump and endothelial pump are: 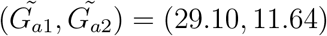. All other parameters are the same for both pumps.

We numerically solve the equations and analyze the spatial distribution of pressure and osmolarity in the circuit. We are interested in how the external mean osmolarity and ion pump energy input influence the total water flux and the flux across the pump.

Node 1-4 correspond to the glomerulus, the renal tubule, the interstitium and the vein, respectively. In general, the pressure drops from the glomerulus to the vein. However, the pressure in the interstitium (node 3) is higher than that in renal tubule (node 2) and vein (node 4). The pressure in interstitium can be further elevated by increasing the epithelial pump energy input 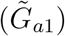, decreasing the blood oncotic pressure (Π*_p_*), or increasing the external osmolarity of permeable solutes (e.g., Π_1_) (Fig. 5 (c)). When varying the osmotic pressure of permeable molecules, the interstitial hydraulic pressure can vary from 5 kPa to 9 kPa (Fig. 5 (e)). In contrast to the hydraulic pressure, the total circuit osmolarity (small molecules and macromolecules) increases monotonically from node 2 to node 4. The osmolarity in the interstitium is in between that of renal tubule and renal vein. The energy input 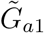 increases the osmotic pressure of the interstitial fluid due to increased ion transport into the interstitium. However, blood oncotic pressure (Π*_p_*) has little effect on the osmotic pressure in the interstitium (Fig. 5 (d)). The interstitial osmolarity can vary up to 0.5 kPa with varying blood oncotic pressure and energy input (Fig. 5 (f)).

Another interesting quantity is the total circulatory water flux. When the mean osmolarity of permeable solutes outside the pump (Π_0_) is increased, both the total water flux and the branch flux across the pump decrease (Fig. S8). As expected, increasing the energy input, 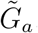, increases the total flux and branch flux (Fig. S8 (a)-(b)). Increasing the blood oncotic pressure (consequently the osmotic pressure difference between node 2 and 4) decreases both the total flux and branch flux across the pump (Fig. S8 (c)-(d)). The branch flux across the pumps can also be determined by locating the intersection between the system curve and the pump performance curve. In the two-pump model, it is not feasible to derive separate systems curves for each pump. Consequently, this section focuses solely on exploring the system curve and pump performance curve corresponding to the epithelial pump. Similar to the one-pump model, an increase in energy input shifts the operating point upward, resulting in larger flux and pressure difference (Fig. 5(g)). However, unlike the one-pump model, external osmolarity changes both the systems curve and pump performance curve (Fig. 5(h)). When active pumping is stopped, both the pressure difference and flux decrease (Fig. 5). Additional results are also given in Fig. S7-S10. It is important to note that, unlike the one-pump circuit, the systems curve here also incorporates information from the “endothelial pump” and the osmotic pressure of permeable solute at node 3, since all elements are coupled together to generate the total circulatory flux.

An interesting prediction in this model is that both the spatial variation and the absolute value of osmotic pressure influences the overall circulation and vessel flux. When the mean external osmotic pressure is increased, water flux decreases. In order to maintain the overall flux at the same level, the pressure from the heart needs to be elevated or blood vessels need to constrict. This can be interpreted as one possible mechanism of hypertension. Another approach for maintaining the water flux is by increasing the energy input and ion pump (e.g. NaK) activity, which can be controlled by hormones and other physiological inputs. Indeed, all parameters in the model are controlled by physiological response, e.g., vessel constriction from smooth muscle action. Therefore, in order to model physiological fluid circulation, additional layer of feedback control is needed. It is likely that hydraulic pressure and osmolarity are constantly sensed by cells in various compartments, and feedback control is used to maintain systemic homeostasis.

## Mechanical efficiency and stress in the pumping element

It is worth noting that for most mechanical pumps, there exists a regime of optimal efficiency and minimal mechanical stress. Similar regimes are likely to exist for cells. If the operating point deviates from this optimal regime, cells are likely to experience increased metabolic and/or physiological stress, potentially predisposing them to various diseases. For example, when hydraulic pressure gradient is altered across the epithelium, cell proteome change was observed [3]. Increased pressure gradient can challenge epithelial integrity and disrupt junctions [30]. We can explore these questions by examining energy efficiency of the pump. In the following discussion, we assume ΔΠ = ΔΠ*_p_* = 0 across the pump. The relative mechanical efficiency can be defined as: 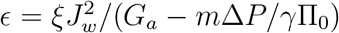, where *ξ* is an effective parameter that includes the surface area of epithelial layer, ATP concentration and ATP hydrolysis rate, etc. With increasing basal-apical pressure difference (Δ*P*), the energy efficiency decreases and eventually reaches 0 at stall pressure Δ*P ^∗^*, where cells experience the maximum mechanical stress. The energy curve also depends on energy input for solute transport 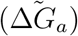 and ion transporter relocalization (*m*) (Fig. 6(a)). In order to achieve optimal energy efficiency and alleviate mechanical stress, cells may potentially adjust the pumping parameters (e.g., water permeability *α*) to modulate the pump performance curve. Additionally, the system curve could be altered in response to signals from stressed cells (Fig. 6(b)). These possible adaptive mechanisms collectively maintain optimal energy efficiency and minimal mechanical stress. If this control system is disrupted, such as mutations observed in PKD [31], diseases may result.

**Figure 6.**
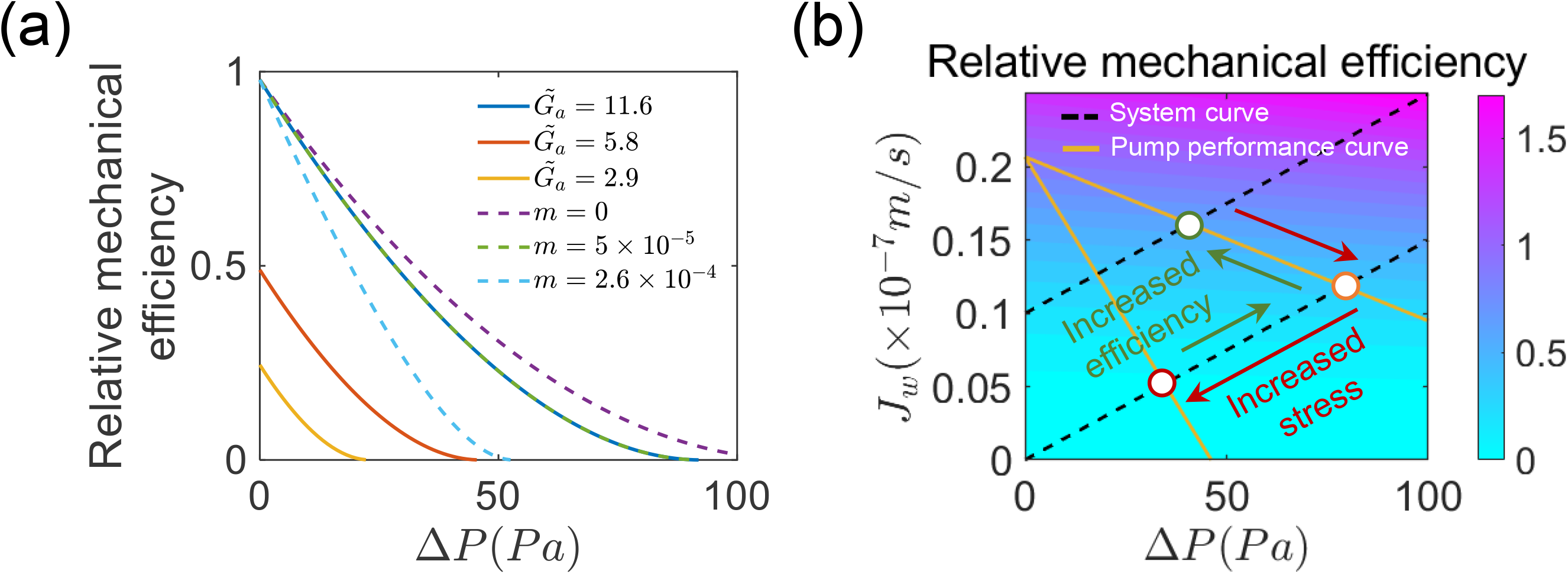
(a) Relative mechanical efficiency as a function of energy input 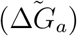 and pressure dependence of solute flux (*m*). (b) Pump performance curves and energy efficiency. Cells could potentially modify both the PPC and the systems curve to enhance mechanical efficiency.

## Discussion and Conclusion

In this paper, we develop a simple physical model explaining the ”pumping” property of renal epithelial and endothelial cells. The model incorporates solute pumping and diffusion, and demonstrates that water transport is facilitated by the establishment of a solute concentration gradient in the cytoplasm. We start with the solute convection-diffusion equations. The equations are solved together with equations of water flux. Our model shows that the cytoplasmic solute concentration profile can follow the external osmolarity change. To ensure normal water flow direction (from apical to basal), the cell actively transports solute so that the inside concentration is higher on the apical side and lower on the basal side relative to the external osmolarity.

Our model allows us to explore the influences of cell energy input on the generalized ”pump performance surface” (water flux vs. pressure gradient and osmolarity gradient) of both water and solute. As expected, the water flux decreases with basal-apical pressure difference and increases with the increase of basal-apical osmotic pressure difference ΔΠ; the solute flux *J_s_* decreases with both ΔΠ and Δ*P*. Interestingly, the absolute value of the external osmolarity also influences the water and solute flux. There are two different regimes where the absolute osmolarity has opposite influences on the flux. The boundary is a function of both pressure and osmolarity gradient. The energy input for ion pump also increases the water and solute flux. We also predict how the osmolarity gradient and the total osmolarity influence the stall pressure for both water and solute. We are able to obtain a phase diagram of water and solute flux, showing regimes of normal pumping and “reversed pumping”. We find that when the energy input goes to zero, we obtain the Starling relation for the permeabilities of epithelia and endothelia.

The cell pumping model serves as the starting point for a realistic physiological circulation model. Previous models of systemic circulation only consider fluid fluxes alone while neglecting solute effects. The novel feature of our work is that the overall fluid circulation is coupled to solute circulation. This allows us to compute systemic properties such as the systems curve (flux vs. pressure gradient in a systematic view) and pressure/osmolarity at various points in the network. When the systems curve is combined together with the pump performance curve, we find the operating point of the pumping element. Note that for most mechanical pumps, there is a regime of highest efficiency/lowest mechanical stress. Similar regime is likely to exist for cells. If the operating point is outside of this regime, cells are likely to experience stress, leading to possible diseases. We also explored how blood oncotic pressure and pump energy input influence the pressure distribution and water flux. The one-pump model predicts that both the total flux and branch flux across the pump decrease with the external osmolarity. Results on how blood oncotic pressure influences the flux provides some understanding on some kidney disease related to protein filtration [32, 33].

A more realistic physiological model consists of two pumps and an interstitium. Again, water and solute circuits are solved together. This model shows a non-monotonic change in pressure across the network - the interstitial pressure is higher than that in the renal tubule and renal vein. The interstitial pressure is influenced by pump energy input, blood oncotic pressure and total blood osmolarity. The predicted osmolarity monotonically increases from the renal tubule to the vein and the osmolarity in interstitial fluid is increased with the increase of ion pump energy input.

Similar to the one-pump model, the two-pump circulation model predicts that both the total flux and branch flux across the pump decrease with the external osmolarity. This result provides a potential explanation of hypertension. Since the circulation flux decreases with increasing osmolarity, the body has to increase the blood pressure to maintain the same circulation flux. Note that our model does not contain possible physiological control of all circulation parameters. In reality, there are multiple ways that pressure, osmolarity, and flux are sensed. Hormone and neural signals can change all parameters in the circuit. The active control of overall circulation is not modeled here. But our work provides a starting point where mechanisms of active control can be assessed.

Results from the two-pump circuit model is different from the one-pump model. This demonstrates that properties of the circulatory network have a significant impact on the behavior of cells. For tissues placed in the system, the pump performance should be solved together with other system parameters. The model can be made more realistic if we can include details of different ionic species, in addition to Na^+^ and Cl*^−^*, other important osmolytes are 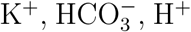 etc. Inclusion of electric fields in cells and tissues [34, 35, 36], which are known to be important for tissue mechanics and morphogenesis, will also expand the range of predicted phenomena. We note that this coupling between cells and the overall circulation system is present in mature organisms as well as during embryo and organ development. Since the overall circulation network influences cell behavior, it is likely that overall circulation can influence cell phenotype specification.

Finally, our current model assumes fixed cells and tissue geometry. Pressure and osmolarity gradients exists across various compartments in the circulation network. This means that there is mechanical stress across the compartment boundaries, which could also impact cell behavior. Pressure gradients will generate deformation and tensions in compartment walls. It is known that cell proliferation and death are influenced by mechanical forces [37, 38]. Moreover, if the compartment walls and boundaries are allowed to deform, and cells are allowed to proliferate and differentiation, then we will arrive at a model of morphogenesis where growth of organs and tissues are coupled to cell mechanical tension/stress, pressure gradients and fluid flux [39]. Indeed, a concrete (beyond phenomenological) model of morphogenesis should include active fluid, nutrient, and O_2_ circulation, as well as the impact of mechanical stress on cell behavior. Our model provides a starting point to examine morphogenesis and development in realistic physiological contexts.

## Author Contributions

SXS, YFW and MAB designed the research. YFW carried out all simulations, analyzed the data. YFW and SXS wrote the article.

## Acknowledgments

This work has been funded in part by R01GM134542.

## References

[1] Cereijido M, Robbins ES, Dolan WJ, Rotunno CA, Sabatini DD. Polarized monolayers formed by epithelial cells on a permeable and translucent support. J. Cell Biol. 1978; 77(3):853–80.

[2] Tripathi S, Boulpaep EL. Mechanisms of water transport by epithelial cells. Q. J. Exp. Physiol. 1989; 74(4):385–417.

[3] Choudhury MI, Li Y, Mistriotis P, Vasconcelos AC, Dixon EE, Yang J, Benson M, Maity D, Walker R, Martin L, Koroma F. Kidney epithelial cells are active mechano-biological fluid pumps. Nat. Commun. 2022; 13(1):2317.

[4] Capurro C, Escobar E, Ibarra C, Porta M, Parisi M. Water permeability in different epithelial barriers. Biol. Cell 1989;66(1-2):145–8.

[5] Wanitschke R, Nell G, Rummel W. Influence of hydrostatic pressure gradients on net transfer of sodium and water across isolated rat colonic mucosa. Naunyn Schmiedebergs Arch. Pharmacol. 1977; 297:191–4.

[6] Misfeldt DS, Hamamoto ST, Pitelka DR. Transepithelial transport in cell culture. Proc. Natl. Acad. Sci. U.S.A. 1976; 73(4):1212–6.

[7] Grandchamp A, Boulpaep EL. Pressure control of sodium reabsorption and intercellular backflux across proximal kidney tubule. J. Clin. Investig. 1974; 54(1):69–82.

[8] Fischbarg J, Warshavsky CR, Lim JJ. Pathways for hydraulically and osmotically-induced water flows across epithelia. Nature. 1977; 266(5597):71–4.

[9] Granger JP, Alexander BT, Llinas M. Mechanisms of pressure natriuresis. Curr. Hypertens. Rep. 2002; 4(2):152–9.

[10] Ivy JR, Bailey MA. Pressure natriuresis and the renal control of arterial blood pressure. J. Physiol. 2014; 592(18):3955–67.

[11] Noordergraaf A. Circulatory system dynamics. Elsevier; 2012.

[12] Titze J. Interstitial fluid homeostasis and pressure: news from the black box. Kidney Int. 2013; 84(5):869–71.

[13] Shieh AC, Swartz MA. Regulation of tumor invasion by interstitial fluid flow. Phys. Biol. 2011; 8(1):015012.

[14] Ruch TC, Patton HD. Physiology and Biophysics, W.B. Saunders; 1976.

[15] Jiang H, Sun SX. Cellular pressure and volume regulation and implications for cell mechanics. Biophys. J. 2013; 105(3):609–19.

[16] Levick JR. Revision of the Starling principle: new views of tissue fluid balance. J. Physiol. 2004; 557(Pt 3):704.

[17] Ferrari E, Palma C, Vesentini S, Occhetta P, Rasponi M. Integrating biosensors in organs-on-chip devices: A perspective on current strategies to monitor microphysiological systems. Biosensors. 2020; 10(9):110.

[18] Wang K, Man K, Liu J, Liu Y, Chen Q, Zhou Y, Yang Y. Microphysiological systems: design, fabrication, and applications. ACS Biomater. Sci. Eng. 2020; 6(6):3231–57.

[19] Kang SM. Recent advances in microfluidic-based microphysiological systems. Biochip J. 2022; 16(1):13–26.

[20] Kung E, Baretta A, Baker C, Arbia G, Biglino G, Corsini C, Schievano S, Vignon-Clementel IE, Dubini G, Pennati G, Taylor A. Predictive modeling of the virtual Hemi-Fontan operation for second stage single ventricle palliation: two patient-specific cases. J. Biomech. 2013; 46(2):423–9.

[21] Kung E, Pennati G, Migliavacca F, Hsia TY, Figliola R, Marsden A, Giardini A, MOCHA Investigators. A simulation protocol for exercise physiology in Fontan patients using a closed loop lumped-parameter model. J. Biomech. Eng. 2014; 136(8):081007.

[22] Migliavacca F, Pennati G, Dubini G, Fumero R, Pietrabissa R, Urcelay G, Bove EL, Hsia TY, de Leval MR. Modeling of the Norwood circulation: effects of shunt size, vascular resistances, and heart rate. Am. J. Physiol. Heart Circ. Physiol. 2001; 280(5):H2076–86.

[23] Snyder MF, Rideout VC. Computer simulation studies of the venous circulation. IEEE. Trans. Biomed. Eng. 1969: 325–34.

[24] Westerhof N, Lankhaar JW, Westerhof BE. The arterial windkessel. Med. Biol. Eng. Comput. 2009; 47(2):131–41.

[25] Feher JJ. Quantitative human physiology: an introduction. Academic press. 2017.

[26] Weinstein AM. A mathematical model of rat distal convoluted tubule. I. Cotransporter function in early DCT. Am. J. Physiol. Renal Physiol. 2005; 289(4):F699–720.

[27] Weinstein AM, Weinbaum S, Duan Y, Du Z, Yan Q, Wong T. Flow-dependent transport in a mathematical model of rat proximal tubule. Am. J. Physiol. 2007; 292.

[28] Bonanno JA. Molecular mechanisms underlying the corneal endothelial pump. Exp. Eye Res. 2012; 95(1):2–7.

[29] Klyce SD. Endothelial pump and barrier function. Exp. Eye Res. 2020; 198:108068.

[30] Duan Y, Gotoh N, Yan Q, Du Z, Weinstein AM, Wang T, Weinbaum S. Shear-induced reorganization of renal proximal tubule cell actin cytoskeleton and apical junctional complexes. Proc. Natl. Acad. Sci. U. S. A. 2008; 105(32):11418–23.

[31] Torres VE, Harris PC. Mechanisms of disease: autosomal dominant and recessive polycystic kidney diseases. Nat. Rev. Nephrol. 2006; 2(1):40–55.

[32] Tryggvason K, Pettersson E. Causes and consequences of proteinuria: the kidney filtration barrier and progressive renal failure. J. Intern. Med. 2003; 254(3):216–24.

[33] Levey AS, Becker C, Inker LA. Glomerular filtration rate and albuminuria for detection and staging of acute and chronic kidney disease in adults: a systematic review. Jama. 2015; 313(8):837–46.

[34] Layton AT, Layton HE. A computational model of epithelial solute and water transport along a human nephron. PLoS Comput. Biol. 2019; 15(2):e1006108.

[35] Yellin F, Li Y, Sreenivasan VK, Farrell B, Johny MB, Yue D, Sun SX. Electromechanics and volume dynamics in nonexcitable tissue cells. Biophys. J. 2018; 114(9):2231–42.

[36] Li Y, Sun SX. The influence of polarized membrane ion carriers and extracellular electrical/pH gradients on cell ionic homeostasis and locomotion. bioRxiv. 2023:2023–07.

[37] Cheng G, Tse J, Jain RK, Munn LL. Micro-environmental mechanical stress controls tumor spheroid size and morphology by suppressing proliferation and inducing apoptosis in cancer cells. PLoS one. 2009; 4(2):e4632.

[38] Zeng Y, Du X, Yao X, Qiu Y, Jiang W, Shen J, Li L, Liu X. Mechanism of cell death of endothelial cells regulated by mechanical forces. J. Biomech. 2022; 131:110917.

[39] Gudipaty SA, Lindblom J, Loftus PD, Redd MJ, Edes K, Davey CF, Krishnegowda V, Rosenblatt J. Mechanical stretch triggers rapid epithelial cell division through Piezo1. Nature. 2017; 543(7643):118–21.

