## Supplementary Note for "Cell-Driven Fluid Dynamics: A Physical Model of Active Systemic Circulation"

### Supplementary Information for "Fluid and Solute Transport by Cells and a Model of Systemic Circulation"

#### Supplementary Note 1: Cell monolayer fluid and solute pumping model

In this section we develop a 1D physical model of the cell monolayer incorporating active solute (e.g., NaCl) transport and demonstrate how hydraulic pressure and osmotic pressure cooperatively contribute to fluid and solute pumping. Here we take the epithelial cells in renal tubule as an example. The cell monolayer can be modeled as a 1D layer. We define the apical (lumen) side of kidney epithelial cells as $x = -L$ and the basal side as $x = L$ (Fig. 1(a), 1(a)). The water flux across the two boundaries can then be expressed as [1]:

$$J_{wa} = \alpha_a[(P_a - P_a^i) - (\Pi_a + \Pi_{p,a} - \Pi_a^i - \Pi_{p,a}^i)] \tag{1}$$

$$J_{wb} = \alpha_b[(P_b^i - P_b) - (\Pi_b^i + \Pi_{p,b}^i - \Pi_b - \Pi_{p,b}^i)] \tag{2}$$

$J_{wa}$ and $J_{wb}$ are water flux (in the unit of m/s) at $x = -L$ (apical side) and $x = L$ (basal side). The directions of water transport are both defined as the positive direction of the x-axis, which points from apical to basal side. $\alpha_a$ and $\alpha_b$ are water permeability at the two boundaries, respectively. $P_a^i, P_b^i$ are the hydrostatic pressure at two boundaries inside the pumping element and $\Pi_a^i, \Pi_b^i$ are the corresponding osmotic pressures from transported small molecules such as NaCl. $\Pi_{pa,pb}^i$. $\Pi_{pa,pb}$ are osmotic pressures from macromolecules such as protein. In the cytoplasm, $\Pi_{p,a}^i = \Pi_{p,b}^i$. For simplicity, we assume that water permeability constants are the same at both ends ($\alpha_a = \alpha_b = \alpha$). At steady state, the water flux across the two boundaries of the pumping element should be equal and the overall water flux across the pumping element can be calculated as:

$$J_w = \frac{1}{2}(J_{wa} + J_{wb}) = \frac{\alpha}{2}\left[-(P_b - P_a) + (P_b^i - P_a^i) + (\Pi_b - \Pi_a) + (\Pi_{p,b} - \Pi_{p,a}) - (\Pi_b^i - \Pi_a^i)\right] \quad (3)$$

If cells in the monolayer are modeled as cylinders, and the internal pressure gradient is given by:

$$J_w = \frac{R_{cell}^2}{8\mu L}(P_a^i - P_b^i) \quad (4)$$

where $R_{cell}$ is the "equivalent radius" and $\mu$ is the dynamic viscosity of the fluid. Note that other assumptions about cell shape does not change the relationship between $J_w$, $\mu$, and pressure difference. Only the prefactor is altered.

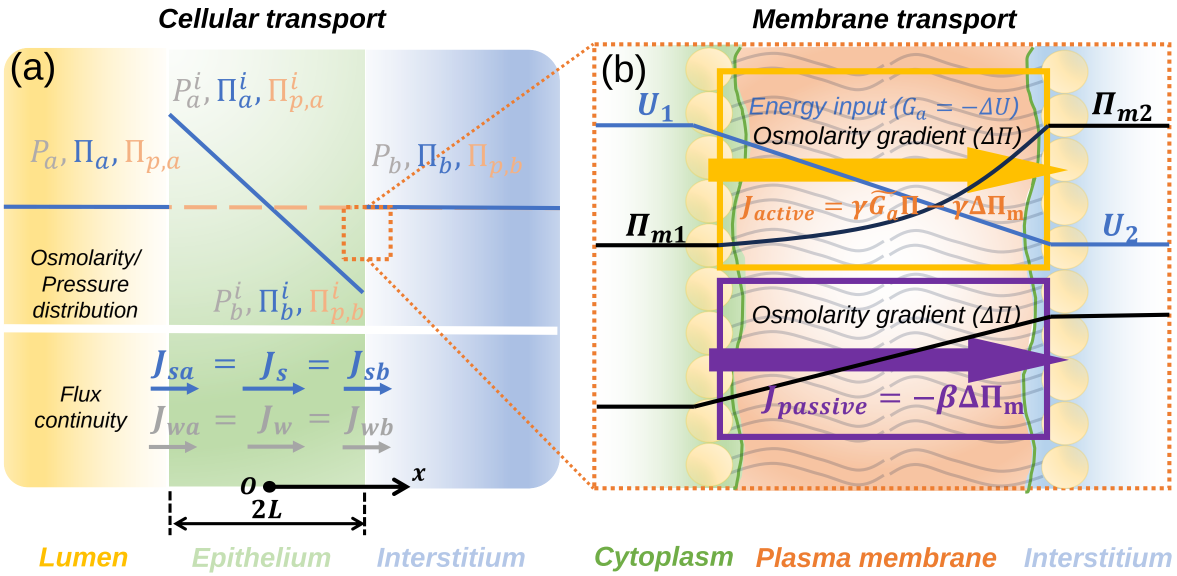


Figure 1: Mathematical illustration of the pump model. (a) Pressure and osmolarity distribution across the cell monolayer. Pressure and osmolarity gradients drive the solute and water fluxes, which are continuous across the cell. (b) Active and passive solute fluxes across the membrane are derived from free energy gradients across the membrane.

To transport water, cells/tissues actively pump ions across the membrane. Consequently, there is a solute concentration gradient inside the cells. For simplicity, we assume there is only one type of solute (e.g., NaCl) involved in the transport process. Moreover, the solute concentration must be continuous in the pumping element and the concentration is described by the convection-diffusion equation. Ion transport occurs at the two ends of the pumping element (Fig. 1(a)). Inside the "pump", the osmotic pressure is:

$$\frac{\partial \Pi}{\partial t} = D\nabla^2\Pi - J_w\nabla\Pi \tag{5}$$

Where $J_w$ is the water flux. For convenience, we replace the concentration by osmotic pressure (only with a coefficient difference given by $\Pi = RTc$). At steady state, the total solute number in the pumping unit should be constant and the flux should be the same everywhere inside the pump, which is given by: $J_s = -D\frac{\partial \Pi}{\partial x} + J_w\Pi$. The solution of the osmotic pressure is then given by: $\Pi = Ce^{\frac{J_w}{D}x} + J_s/J_w$, where $C$ is a constant determined by the boundary condition. Since $J_w$ is around $1.6 \times 10^{-7}$m/s [2] and the diffusion coefficient $D$ of ions is in the order of $10^{-9}\mathrm{m}^2/\mathrm{s}$ [3, 4], we naturally have $\frac{J_w}{D}x \sim 10^{-3} << 1$. Here we take the length scale $x = 10\mu m$, the same as a cell. Given the small exponent, we can approximate the osmotic pressure profile inside the cell as

$$\Pi = \frac{J_w C}{D}x + \left(C + \frac{J_s}{J_w}\right). \tag{6}$$

To obtain the constant $C$, we need to know the solute flux at the cell apical and basal boundaries, which can be obtained from the following. There are two types of ion transporters on the membrane: ion pumps and ion channels (Fig. 1(b)). For the ion pump, the ion transport is accompanied by energy input, and the transport direction can be against the concentration gradient. In general, the velocity of an ion through the membrane can be expressed as: $\boldsymbol{v_a} = -\gamma'\nabla\mu$, where $\gamma'$ is a proportionality constant. $\mu = RT\ln c + U$ is the chemical potential of noninteracting particles of concentration $c$. $R$ is gas constant and $T$ is temperature. $U$ is the driving potential corresponding to the energy input (e.g., energy from ATP hydrolysis). The solute flux in the membrane is then:

$$J_a = -\gamma' c\nabla(RT\ln c + U) \tag{7}$$

In 1D, if we assume that $U$ is a linear function of $x$, $\nabla U = (U_2 - U_1)/h = -G/h$. Here $G$ is the potential difference across the membrane and $h$ is the membrane thickness. Eq. 7 can then be rewritten as: $J_a = -\frac{\gamma' RT}{h}\left(h\frac{\partial c}{\partial x} - c\tilde{G}\right)$. Here $\tilde{G} = G/RT$ is the dimensionless energy input. Multiplying both sides by $RT$, we can write the solute flux in terms of osmotic pressure in the membrane as:

$$J_a = \gamma\left(\Pi\tilde{G} - h\frac{\partial \Pi}{\partial x}\right) \tag{8}$$

Where $\gamma = \frac{\gamma' RT}{h}$. At steady state, the solute flux $J_a$ should be constant inside the membrane. Given a constant $J_a$, eq. 8 can then be solved as: $\Pi = C' e^{\frac{\tilde{G}}{h}x} + \frac{J_a}{\gamma \tilde{G}}$, where $C'$ is an integration constant. Given the osmolarity at both sides of the membrane (Fig. 1(b)): $\Pi|_{x=0} = \Pi_{m1}$ and $\Pi|_{x=h} = \Pi_{m2}$, we can determine the constant $C'$ and the final expression for ion flux through the pump as:

$$J_a = \gamma \tilde{G} \left( \Pi_{m1} - \frac{\Delta \Pi_m}{e^{\tilde{G}} - 1} \right) \tag{9}$$

where $\Delta \Pi_m = \Pi_{m2} - \Pi_{m1}$ is the osmolarity difference across the cell membrane.

Similarly, we can obtain the solute flux across the ion channels by setting external potential $U = 0$. The passive ion flux is then: $J_p = -\beta \Delta \Pi_m$. Notably, inside the ion pump, the osmolarity is exponentially distributed, while inside the ion channel, the osmolarity is linear (Fig. 1(b)).

Biologically, the energy input is related to the activity state and the number ion pumps in the membrane. In experiments, it was found that there are polarization mechanisms that can alter ion pump localization, which means that the energy input can depend on the external hydrostatic pressure and osmotic pressure [5, 6].In general, the energy input is a complex function of the form $\tilde{G} = h(\tilde{G_a}, \frac{\Delta P}{\Pi_{m1}}, \frac{\Delta\Pi + \Delta\Pi_p}{\Pi_{m1}})$. To simplify, we can approximate to linear order: $\tilde{G} \approx \tilde{G_a} - m\frac{\Delta P}{\Pi_{m1}} - m' \frac{\Delta\Pi + \Delta\Pi_p}{\Pi_{m1}}$. Here $\Delta P = P_b - P_a$, $\Delta\Pi = \Pi_b - \Pi_a$ and $\Delta\Pi_p = \Pi_{p,b} - \Pi_{p,a}$ correspond to the gradient across the cell (Fig. 1(a)). The coefficients $m, m'$ describe how pressure and osmolarity can influence the localization of ion pumps in response to the pressure and osmolarity gradient. Here, in accordance with experimental observation, we assume that both the osmolarity gradient and pressure gradient decrease the ion pump density thus decreasing the solute flux.

Let $\frac{\tilde{G}}{e^{\tilde{G}}-1} = g(\tilde{G_a}, \Delta P/\Pi_{m1}, (\Delta\Pi + \Delta\Pi_p)/\Pi_{m1})$. Since $\Delta P/\Pi_{m1} \ll 1, (\Delta\Pi + \Delta\Pi_p)/\Pi_{m1} \ll 1$, eq. 9 is approximately:

$$J_a \approx \gamma[\tilde{G_a}\Pi_{m1} - m\Delta P - m'(\Delta\Pi + \Delta\Pi_p) - g(\tilde{G_a}, 0, 0)\Delta\Pi_m] \tag{10}$$

Here, $g(\tilde{G_a}, \Delta P/\Pi_{m1}, (\Delta\Pi + \Delta\Pi_p)/\Pi_{m1})$ is linearly expanded and all the second order terms of $\Delta P/\Pi_1$, $\Delta\Pi/\Pi_1$, $\Delta\Pi_m/\Pi_1$ are dropped. Adding the flux through the pump and the channel together, we obtain the total flux through the membrane:

$$J_s = J_a + J_p = \gamma\tilde{G_a}\Pi_{m1} - \eta\Delta\Pi_m - m\Delta P - m'(\Delta\Pi + \Delta\Pi_p) \tag{11}$$

Where $\eta = [\gamma g(\tilde{G_a}, 0, 0) + \beta]$. Typically $\tilde{G_a}$ takes the value around 10 and $\gamma g(\tilde{G_a}, 0, 0) << \beta$, therefore $\eta \approx \beta$. Here we incorporate $\gamma$ into the coefficients $m$ and $m'$ for simplicity.

The length of the cell (pumping element) is $2L$ and the coordinate origin is placed at the center of the pumping unit: The flux through the apical and basal membranes at $x = -L$ (renal tubule) and $x = L$ (vein) can then be written using Eq. 11:

$$J_s|_{x=-L} = \gamma \tilde{G_{a1}} \Pi_a - \eta(C - \frac{CJ_w}{D}L + \frac{J_s}{J_w} - \Pi_a) - m\Delta P - m'(\Delta\Pi + \Delta\Pi_p) \tag{12}$$

$$J_s|_{x=L} = \gamma \tilde{G_{a2}} \Pi_b^i + \eta(C + \frac{CJ_w}{D}L + \frac{J_s}{J_w} - \Pi_b) - m\Delta P - m'(\Delta\Pi + \Delta\Pi_p) \tag{13}$$

Here $\Delta P = P_b - P_a$. $\Pi_b^i = \frac{CJ_wL}{D} + \frac{J_s}{J_w} + C$ is the osmotic pressure inside the pumping element at the basal side ($x = L$). The energy input at the apical/basal sides, $\tilde{G_{a1}}$ and $\tilde{G_{a2}}$, can be different, but for simplicity, we assume $\tilde{G_{a1}} = \tilde{G_{a2}} = \tilde{G_a}$. The coefficients $m$ and $m'$ describe the solute flux dependence on pressure and osmolarity, which include the influence of pressure and osmolarity on ion pump distribution and energy input for ion pumping.

Combing eqs 3,4, 12 and 13, we can obtain the three unknowns: solute ($J_s$) and water flux ($J_w$) across the pumping element, and $C$. Also, we can obtain the osmolarity profile in the pumping element. The solution is as follows: To begin, we define: $x_1 = C + J_s/J_w, x_2 = CJ_wL/D, x_3 = CL/D$. The solute equations 12 and 13 can then be simplified as:

$$\frac{x_1x_2}{x_3} - \frac{Dx_2}{L} = \eta r\Pi_a - \eta(x_1 - x_2 - \Pi_a) - m\Delta P - m'(\Delta\Pi + \Delta\Pi_p) \tag{14}$$

$$\frac{x_1x_2}{x_3} - \frac{Dx_2}{L} = \eta(1 + r)(x_1 + x_2) - \eta\Pi_b - m\Delta P - m'(\Delta\Pi + \Delta\Pi_p) \tag{15}$$

Where $r = \frac{\gamma}{\eta}\tilde{G_a}$ is dimensionless energy input. Substituting eq. 4 into 3, we can remove the unknown $(P_a^i - P_b^i)$:

$$\left(\frac{8\mu L\alpha}{R_{cell}^2} + 2\right) J_w = \alpha(-\Delta P + \Delta\Pi + \Delta\Pi_p - 2x_2) \tag{16}$$

Let $\alpha_s = \frac{\alpha}{8\mu L\alpha/R_{cell}^2 + 2}$, the equation above can be simplified as:

$$J_w = \frac{x_2}{x_3} = \alpha_s(-\Delta P + \Delta\Pi + \Delta\Pi_p - 2x_2) \tag{17}$$

The solution is given by eqs. 14, 15 and 17. From eqs 14 and 15, we obtain:

$$x_1 + \theta x_2 = \Pi_0 \tag{18}$$

In eq. 18, $\theta = r/(r+2)$ and $\Pi_0 = \frac{(r+1)\Pi_a + \Pi_b}{r+2}$ is a weighted mean osmotic pressure. This weighted mean osmotic pressure incorporates both apical and basal osmotic pressure, and comes from the continuity of the solute flux $J_s$ across the monolayer. Substituting eqs. 17 and 18 into eq. 14, we can eliminate variables $x_1$ and $x_3$ and finally obtain a quadratic equation of $x_2$:

$$2\theta\alpha_s x_2^2 - [2\alpha_s\Pi_0 + \alpha_s\theta(-\Delta P + \Delta\Pi + \Delta\Pi_p) + \frac{D}{L} + \eta(\theta+1)]x_2 + [\alpha_s\Pi_0(-\Delta P + \Delta\Pi + \Delta\Pi_p) - \eta\Pi_{02} + m\Delta P + m'(\Delta\Pi + \Delta\Pi_p)] = 0 \quad (19)$$

In eq. 19, $\Pi_{02} = \frac{(r+1)^2\Pi_a - \Pi_b}{r+2}$. Eq. 19 is a quadratic equation of $x_2$, and there are two roots. In order to get positive water flux with realistic values, we take the root:

$$x_2 = \frac{B - \sqrt{B^2 - 4AC_2}}{2A} \quad (20)$$

Other unknowns can be solved as:

$$x_1 = \Pi_0 - \theta x_2 \quad (21)$$

$$x_3 = \frac{x_2}{\alpha_s(-\Delta P + \Delta\Pi + \Delta\Pi_p) - 2\alpha_s x_2} \quad (22)$$

In eqs. 20 - 22, $A = 2\theta\alpha_s, B = 2\alpha_s\Pi_0 + \alpha_s\theta(-\Delta P + \Delta\Pi + \Delta\Pi_p) + \frac{D}{L} + \eta(\theta+1), C_2 = \alpha_s\Pi_0(-\Delta P + \Delta\Pi + \Delta\Pi_p) - \eta\Pi_{02} + m\Delta P + m'(\Delta\Pi + \Delta\Pi_p)$. $\Delta P = P_b - P_a$, $\Delta\Pi = \Pi_b - \Pi_a$ and $\Delta\Pi_p = \Pi_{p,b} - \Pi_{p,a}$. With $x_1 \sim x_3$, we are able to obtain water and solute fluxes from eqs. 14 and 17.

From eq. 14, 15, 17, we see that the water transport rate is closely coupled with the solute transport. These two processes are determined by both external variables (e.g. osmolarity gradient ($\Delta\Pi$), pressure gradient ($\Delta P$) and absolute osmolarity ($\Pi_0$)) and internal variables (e.g., water permeability of the membrane ($\alpha$) and energy input for active ion pumping ($\tilde{G_a}$)). It's the coupled fields of osmolarity and pressure that determine the fluid transport through the pumping element.

The exact solution is not physically intuitive and it is hard to interpret analytically. However, we can further simplify the results using scaling analysis. Since $AC_2/B^2 \sim 10^{-8} \ll 1$, eq. 20 is:

$$x_2 \approx C_2/B = \frac{\alpha_s \Pi_0(-\Delta P + \Delta\Pi + \Delta\Pi_p) + m\Delta P + m'(\Delta\Pi + \Delta\Pi_p) - \eta\Pi_{02}}{2\alpha_s\Pi_0 + \alpha_s\theta(-\Delta P + \Delta\Pi + \Delta\Pi_p) + \eta(\theta+1) + D/L} \tag{23}$$

We can further simplify the solution by comparing the order of magnitude for each term. Since $\Delta P \sim 100Pa$ and $\Delta\Pi = \Delta\Pi_p \approx 3.8 \times 10^3$ Pa, the scale of each term is: $\alpha_s\Pi_0(-\Delta P + \Delta\Pi + \Delta\Pi_p) \approx 3.01, \eta\Pi_{02} \approx -0.47, 2\alpha_s\Pi_0 \approx 8\times10^{-4}, \alpha_s\theta(-\Delta P + \Delta\Pi + \Delta\Pi_p) \approx 2.35\times10^{-10}, \eta(\theta+1) \approx 2.6\times10^{-4}, D/L \approx 2\times10^{-4}$. Also, since $\theta \approx 6.27\times10^{-5} \ll 1$, $\eta(\theta+1) \approx \eta$. The coefficients $m, m'$ can vary in different situations. Since we do not have an accurate value for $m\Delta P$ and $m'\Delta\Pi$, we assume that these two terms are in the same order as the biggest term and therefore the pressure-dependent term and osmolarity-dependent term are always kept. After dropping small terms, we obtain the first approximation:

$$x_2 = \frac{\alpha_s \Pi_0(-\Delta P + \Delta\Pi + \Delta\Pi_p) + m\Delta P + m'(\Delta\Pi + \Delta\Pi_p) - \eta\Pi_{02}}{2\alpha_s\Pi_0 + D/L + \eta} \tag{24}$$

Further approximations can be done given that $\Delta\Pi \ll \Pi_a \approx \Pi_b$. The apical and basal osmotic pressure can be approximated by the mean osmotic pressure $\Pi_0$: $\Pi_a \approx \Pi_0, \Pi_b \approx \Pi_0 + \Delta\Pi$. With this approximation, $\Pi_{02} \approx \frac{(r+1)^2\Pi_0 - \Pi_0 - \Delta\Pi}{r+2} = r\Pi_0 - \frac{1}{r+2}\Delta\Pi$. Further, according to eq. 17, the water flux can be approximated as:

$$J_w = \frac{\alpha_s(D+\eta L+2mL)}{2\alpha_s\Pi_0 L + (D+\eta L)}\left(-\Delta P + \frac{D+\theta\eta L - 2m'L}{D+\eta L+2mL}\Delta\Pi + \frac{D+\eta L-2m'L}{D+\eta L+2mL}\Delta\Pi_p + \frac{2\eta r L}{D+\eta L+2mL}\Pi_0\right) \tag{25}$$

Eq. 25 can be further simplified by letting $\alpha_{ss} = \frac{\alpha_s(D+\eta L+2mL)}{2\alpha_s\Pi_0 L+(D+\eta L)}, \zeta_{w1} = \frac{D+\theta\eta L-2m'L}{D+\eta L+2mL}, \zeta_{w2} = \frac{D+\eta L-2m'L}{D+\eta L+2mL}, \zeta_{w3} = \frac{2\eta r L}{D+\eta L+2mL}$, which gives the final result for the overall flux through the cell:

$$J_w = \alpha_{ss}(-\Delta P + \zeta_{w1}\Delta\Pi + \zeta_{w2}\Delta\Pi_p + \zeta_{w3}\Pi_0). \tag{26}$$

The flux expression resembles that of a water pump, which is in the form: $J_w = \alpha(\Delta P - \Delta P^*)$. $\Delta P^*$ is the stall pressure corresponding to zero flux. We can see that the stall pressure for the epithelial (or endothelial) pumping element is determined by both osmotic pressure gradient $(\Delta\Pi, \Delta\Pi_p)$ and the mean osmotic pressure in the environment $(\Pi_0)$. Both osmotic pressure difference and mean osmolarity increase the stall pressure. The stall pressure depends on the environmental condition.

Similarly, we obtain an analytical approximation for the solute flux $J_s$ from eq. 14:

$$J_s = \eta[\Pi_{02} + (\theta+1)x_2] - m\Delta P - m'(\Delta\Pi + \Delta\Pi_p) \tag{27}$$

After dropping $\theta$ and substituting $\Pi_{02} \approx \frac{(r+1)^2\Pi_0 - \Pi_0 - \Delta\Pi}{r+2} = r\Pi_0 - \frac{1}{r+2}\Delta\Pi$ into the equation above, we have:

$$J_s = \frac{(\eta+2m)\alpha_s L\Pi_0 + mD}{2\alpha_s\Pi_0 L + D + \eta L}\left\{-\Delta P + \frac{[r(\eta-2m') - 4m']\alpha_s\Pi_0 L - [\eta + (r+2)m']D}{[(\eta+2m)\alpha_s\Pi_0 L + mD](r+2)}\Delta\Pi \right. \\ \left. + \frac{(\eta-2m')\alpha_s\Pi_0 L - m'D}{(\eta+2m)\alpha_s\Pi_0 L + mD}\Delta\Pi_p + \frac{\eta r(2\alpha_s\Pi_0 L + D)}{(\eta+2m)\alpha_s\Pi_0 L + mD}\Pi_0\right\} \tag{28}$$

This can be rewritten as:

$$J_s = \alpha'_{ss}(-\Delta P + \zeta_{s1}\Delta\Pi + \zeta_{s2}\Delta\Pi_p + \zeta_{s3}\Pi_0) \tag{29}$$

Where $\zeta_{s1} = \frac{[r(\eta-2m')-4m']\alpha_s\Pi_0 L - [\eta+(r+2)m']D}{[(\eta+2m)\alpha_s\Pi_0 L + mD](r+2)}, \zeta_{s2} = \frac{(\eta-2m')\alpha_s\Pi_0 L - m'D}{(\eta+2m)\alpha_s\Pi_0 L + mD}, \zeta_{s3} = \frac{\eta r(2\alpha_s\Pi_0 L + D)}{(\eta+2m)\alpha_s\Pi_0 L + mD}$.

In the analysis above, we only considered active transcellular transport process. In reality, there could also be paracellular transport, which refers to the flux passing through the intercellular space between the cell junctions. Paracellular flux is passive, and is directly determined by the pressure difference and osmolarity difference across the cell. The paracellular water and solute transport can be expressed as: $J_{w,para} = \sigma_w(-\Delta P + \Delta\Pi + \Delta\Pi_p), J_{s,para} = -\sigma_s\Delta\Pi$. We can add these terms to the transcellular fluxes to obtain the total flux, which effectively modify the coefficients in eq. 25 and 28.

An interesting result of our analysis is that the absolute external osmolarity also contributes to the water and solute flux. There are two regimes where absolute osmolarity $\Pi_0$ has different influences on water and solute flux depending on osmolarity difference and pressure difference (Fig. 3(a)(b)). The critical condition distinguishing these two regimes can be estimated from the analytical approximation. For water flux, the critical condition is: $(\frac{2m\alpha_s}{\eta+D/L} + \alpha_s)\Delta P + [\frac{2\alpha_s}{\eta+D/L}(m' + \frac{\eta}{r+2}) - \alpha_s]\Delta\Pi + [\frac{2\alpha_s}{\eta+D/L}m' - \alpha_s]\Delta\Pi_p = -\eta r$; For solute flux, the critical condition is: $(\frac{2m\alpha_s}{\eta+D/L} + \alpha_s)\Delta P - [\alpha_s - \frac{2\alpha_s}{\eta+D/L}(m' + \frac{\eta}{r+2})]\Delta\Pi - [\alpha_s - \frac{2\alpha_s}{\eta+D/L}m']\Delta\Pi_p = \frac{r(2\alpha_s\Pi_0 + \eta + D/L)^2}{\eta+D/L} - \eta r$.

Another interesting quantity is stall pressure, which represents the maximum pressure gradient that the cell can sustain with normal water (solute) transport. The approximated stall pressure for water and solute can be obtained from eq. 25, 28:

$$\Delta P_w^* = \frac{D+\theta\eta L-2m'L}{D+\eta L+2mL}\Delta\Pi + \frac{D+\eta L-2m'L}{D+\eta L+2mL}\Delta\Pi_p + \frac{2\eta rL}{D+\eta L+2mL}\Pi_0 \tag{30}$$

$$\Delta P_s^* = \frac{[r(\eta-2m')-4m']\alpha_s\Pi_0 L-[\eta+(r+2)m']D}{[(\eta+2m)\alpha_s\Pi_0 L+mD](r+2)}\Delta\Pi + \frac{\eta r(2\alpha_s\Pi_0 L+D)}{(\eta+2m)\alpha_s\Pi_0 L+mD}\Pi_0 + \frac{(\eta-2m')\alpha_s\Pi_0 L-m'D}{(\eta+2m)\alpha_s\Pi_0 L+mD}\Delta\Pi_p \tag{31}$$

Additional results are presented to illustrate the dependencies of water and solute fluxes on the pressure sensitivity of solute flux ($m$ and $m'$) (Fig. 2), solute and water transport coefficients ($\gamma, \eta, \alpha$) (Fig. 3), and osmotic pressure from impermeable macromolecules (Fig. 4).

### Supplementary Note 2: A systemic Model of Fluid Circulation

#### A one-pump circuit model

We firstly develop a one-pump circuit model in which there is only one pumping element. This pumping element effectively models the epithelium, the interstitium, and the endothelium as a combined unit. The inlet of the pump is the lumen side of the renal tubule and the outlet faces the lumen of blood vessels. A schematic illustration of the model is shown in Fig. 4(a). In general, blood is pumped from the heart (modelled as a source with pressure $P_s$) and goes through aorta ($R_A$). It branches before the kidney: some directly goes through the capillary blood vessel of the organs (effectively modelled as one resistor $R_O$) while others are filtered by the glomerulus ($R_G$) and then goes through the pumping unit consisting of epithelium, interstitium and endothelium (water reabsorption). The two branches finally merge and go back to heart through vena cava ($R_V$). For convenience, we define the node pressure at the end of vena cava to be zero. All the capacitors can be combined together as one effective capacitor $C_E$, which eventually gives a trivial prediction on dynamical response of the pressure and flux. Therefore, we will neglect the "capacitors" and focus on the static property of the system. Similarly, we can define the circuit of osmolytes, in which the hydrostatic pressure is replaced by osmotic pressure. The osmotic pressure remains the same through the blood capillaries in organs but changes across the glomerulus. The plasma proteins do not go through the glomerulus and therefore there is an oncotic pressure difference $\Pi_P$, which is approximately $\Pi_1 - \Pi_2 = \Pi_P = 3.8kPa$. Similar to electrical circuit, we use Kirchhoff's current law to establish the equations for pressure. Different from Ohm's law, the flux is determined by both hydrostatic pressure and osmotic pressure as: $I = (\Delta P - \Delta\Pi)/R$. There are three nodes in the network and the equations are:

$$\frac{P_s - P_1}{R_A} = \frac{P_1 - P_3}{R_O} + \frac{P_1 - P_2 - \Pi_P}{R_G} \tag{32}$$

$$\frac{P_1 - P_2 - \Pi_P}{R_G} = \alpha_{ss}[-(P_3 - P_2) + \Delta P^*]S_{Ep} \tag{33}$$

$$\alpha_{ss}[-(P_3 - P_2) + \Delta P^*]S_{Ep} + \frac{P_1 - P_3}{R_O} = \frac{P_3}{R_V} \tag{34}$$

where $R_A, R_O, R_G, R_V$ are resistances of aorta, capillaries in organs, glomerulus and vena cava, respectively. $P_s$ is the pressure of the heart, and $P_1, P_2, P_3$ are the node pressures. $\Pi_1, \Pi_3$ are osmotic pressure at nodes 1 and 3. Only $(\Pi_1 - \Pi_2) = \Pi_P$ is included in the equations because $\Pi_1 = \Pi_3$ and $\Pi_2$ is the only one different. The pumping performance of the pump is given as: $I = \alpha_{ss}[\Delta P^* - (P_3 - P_2)]$, where $\alpha$ is a permeability constant and $\Delta P^*$ is the "stall pressure" given by eq. 30. The coefficient $S_{Ep}$ is the total surface area of the renal tubule, which transforms the velocity (m/s) into the volume flow rate ($m^3$/s). By defining $R_O = R, R_A = k_1 R, R_G = k_2 R, R_V = k_3 R$, we obtain analytical solutions for all node pressures:

$$P_1 = \frac{[(1 + k_3) + R\alpha_{ss}S_{Ep}(k_2 + k_3 + k_2k_3)]P_s + k_1 R\alpha_{ss}S_{Ep}(\Pi_P - \Delta P^*)}{(1 + k_1 + k_3) + R\alpha_{ss}S_{Ep}(k_1 + k_2 + k_3 + k_1k_2 + k_2k_3)} \tag{35}$$

$$P_2 = \frac{[k_3 + 1 + k_3(k_2 + 1)R\alpha_{ss}S_{Ep}]P_s - (1 + k_1 + k_3 + k_3R\alpha_{ss}S_{Ep})\Pi_P - (k_1 + k_2 + k_1k_2 + k_2k_3)R\alpha_{ss}S_{Ep}\Delta P^*}{(1 + k_1 + k_3) + R\alpha_{ss}S_{Ep}(k_1 + k_2 + k_3 + k_1k_2 + k_2k_3)} \tag{36}$$

$$P_3 = \frac{k_3[(1 + (k_2 + 1)R\alpha_{ss}S_{Ep})P_s + R\alpha_{ss}S_{Ep}(\Delta P^* - \Pi_P)]}{(1 + k_1 + k_3) + R\alpha_{ss}S_{Ep}(k_1 + k_2 + k_3 + k_1k_2 + k_2k_3)} \tag{37}$$

We can also obtain the total flux and branch flux through the pump:

$$I_t = \frac{[1/R + \alpha_{ss}S_{Ep}(k_2 + 1)]P_s - \alpha_{ss}S_{Ep}(\Pi_P - \Delta P^*)}{(1 + k_1 + k_3) + R\alpha_{ss}S_{Ep}(k_1 + k_2 + k_3 + k_1k_2 + k_2k_3)} \tag{38}$$

$$I_{23} = \frac{\alpha_{ss}S_{Ep}[P_s - (1 + k_1 + k_3)(\Pi_P - \Delta P^*)]}{(1 + k_1 + k_3) + R\alpha_{ss}S_{Ep}(k_1 + k_2 + k_3 + k_1k_2 + k_2k_3)} \tag{39}$$

The solutions above are general expressions for systematic quantities since the stall pressure can be any function. In fact, we can use the stall pressure $\Delta P^*$ in Eq. 30. Effects of osmotic pressure on node pressures and fluxes are presented in Fig. 5. Fig. 6 shows how fluxes and pressure distribution are influenced by ion and water transport coefficients in the one-pump model. Note that in the one-pump model, the pumping element consists of epithelial layer, endothelial layer, and interstitium. We set the effective water transport constant as $\alpha = 5 \times 10^{-11}\ m \cdot s^{-1} \cdot Pa^{-1}$ to ensure that the water flux across the pump is in the realistic range ($J_w \sim 10^{-6} m^3/s$).

Another interesting aspect of the circuit is the systems curve, which describes the relation between flux and pressure drop across the pump from a systematic view. The intersection of pump performance curve and systems curve gives the operating parameters of the pump when placed into the circulatory circuit. When placed into the system, the pump may behave differently under the influence of the system. On the other hand, the systemic variables will also be influenced by the pumping element. The systems curve can be directly derived from eqs. 32 - 34. From eq. 32, we obtain the expression of $P_1$ in terms of $P_2$ and $P_3$:

$$P_1 = \frac{k_1}{k_1 + k_2 + k_1 k_2} P_2 + \frac{k_1 k_2}{k_1 + k_2 + k_1 k_2} P_3 + \frac{k_2 P_s + k_1 \Pi_P}{k_1 + k_2 + k_1 k_2} \tag{40}$$

Substituting the equation above into Eqs. 33 and 34, we obtain the branch flux through the pump in terms of only $P_2$ and $P_3$:

$$I_{23} = \frac{1}{R}\left[\frac{k_1}{k_1 + k_2 + k_1 k_2} P_3 - \frac{k_1 + 1}{k_1 + k_2 + k_1 k_2} P_2 + \frac{P_s - (k_1 + 1)\Pi_P}{k_1 + k_2 + k_1 k_2}\right] \tag{41}$$

$$I_{23} = \frac{1}{R}\left[\frac{k_1 + k_2 + k_1 k_2 + k_1 k_3 + k_2 k_3}{k_3(k_1 + k_2 + k_1 k_2)} P_3 - \frac{k_1}{k_1 + k_2 + k_1 k_2} P_2 - \frac{k_2 P_s + k_1 \Pi_P}{k_1 + k_2 + k_1 k_2}\right] \tag{42}$$

Multiply eq. 41 by $(k_1 + k_2 + k_1 k_2 + k_2 k_3)$ and eq. 42 by $k_3$. Adding the two equations together we obtain the systems curve as:

$$I_{23} = \frac{1}{R} \frac{(k_1 + k_3 + 1)(\Delta P - \Pi_P) + P_s}{k_1 + k_2 + k_3 + k_1 k_2 + k_2 k_3}. \tag{43}$$

#### A two-pump circuit model including interstitium

In the previous section, epithelial tissue and endothelial tissue are combined together with interstitium to form an equivalent pump. However, in physiology, there are osmolarity gradients in different compartments established by active ion transport. The differences in osmotic pressure and hydrostatic pressure in the three compartments play an important role in systemic water transport. In this part, we develop a more detailed model which includes the renal epithelial pump, the endothelial pump and the interstitium. Figure 5 (a)-(b) show the structure of the circuit. Node 3 corresponds to the interstitium. Similarly, the two pump model includes both the hydrostatic pressure circuit and the osmotic pressure circuit. According to Kirchhoff's current law, the hydrostatic pressures of the four nodes can be described by:

$$\frac{P_s - P_1}{R_A} = \frac{P_1 - P_4}{R_O} + \frac{P_1 - P_2 - \Pi_P}{R_G} \tag{44}$$

$$\frac{P_1 - P_2 - \Pi_P}{R_G} = \alpha_{ss1}[-(P_3 - P_2) + \Delta P_1^*]S_{Ep} \tag{45}$$

$$\alpha_{ss1}[-(P_3 - P_2) + \Delta P_1^*]S_{Ep} = \alpha_{ss2}[-(P_4 - P_3) + \Delta P_2^*]S_{Ed} \tag{46}$$

$$\alpha_{ss2}[-(P_4 - P_3) + \Delta P_2^*]S_{Ed} + \frac{P_1 - P_4}{R_O} = P_4/R_V \tag{47}$$

where, $\Delta P_1^* = \frac{D+\theta_1\eta_1 L - 2m_1' L}{D+\eta_1 L+2m_1 L}(\Pi_3 - \Pi_2) + \frac{2\eta_1 r_1 L}{D+\eta_1 L+2m_1 L}\Pi_{0p}$, $\Delta P_2^* = \frac{D+\theta_2\eta_2 L-2m_2' L}{D+\eta_2 L+2m_2 L}(\Pi_4 - \Pi_3) + \frac{D+\eta_2 L-2m_2' L}{D+\eta_2 L+2m_2 L}\Pi_p + \frac{2\eta_2 r_2 L}{D+\eta_2 L+2m_2 L}\Pi_{0d}$. In the stall pressures, $\Pi_{0p}$ and $\Pi_{0d}$ are the mean osmotic pressures around renal epithelial pump and endothelial pump, respectively:

$$\Pi_{0p} = \frac{(r_1+1)\Pi_2 + \Pi_3}{r_1+2} \tag{48}$$

$$\Pi_{0d} = \frac{(r_2+1)\Pi_3 + \Pi_4}{r_2+2} \tag{49}$$

This system has 7 unknowns while there are only six equations. In order to close the system, we need another condition, which is provided by the solute flux balance in the interstitium ($J_{s1} = J_{s2}$), which can be obtained from eq. 28:

$$\begin{aligned}\alpha'_{ss,1}[-(P_3-P_2)+\frac{[r_1(\eta_1 - 2m_1') - 4m_1']\alpha_{s1}\Pi_{0p}L - [\eta_1 + (r_1+2)m_1']D}{[(\eta_1+2m_1)\alpha_{s1}\Pi_{0p}L + m_1 D](r_1+2)}(\Pi_3-\Pi_2)+\frac{\eta_1 r_1(2\alpha_{s1}\Pi_{0p}L + D)}{(\eta_1+2m_1)\alpha_{s1}\Pi_{0p}L + m_1 D}\Pi_{0p}]S_{Ep} = \\ \alpha'_{ss,2}[-(P_4-P_3) + \frac{[r_2(\eta_2 - 2m_2') - 4m_2']\alpha_{s2}\Pi_{0d}L - [\eta_2 + (r_2+2)m_2']D}{[(\eta_2+2m_2)\alpha_{s2}\Pi_{0d}L + m_2 D](r_2+2)}(\Pi_4 - \Pi_3) + \frac{\eta_2 r_2(2\alpha_{s2}\Pi_{0d}L + D)}{(\eta_2+2m_2)\alpha_{s2}\Pi_{0d}L + m_2 D}\Pi_{0d} + \\ \frac{(\eta_2 - 2m_2')\alpha_{s2}\Pi_{0d}L - m_2' D}{(\eta_2+2m_2)\alpha_{s2}\Pi_{0d}L + m_2 D}\Pi_p]S_{Ed}\end{aligned} \tag{50}$$

In equations above: subscripts 1 and 2 corresponds to epithelial and endothelial pumps respectively. The solute flux coefficients are $\alpha'_{ss,i} = \frac{(\eta_i+2m_i)\alpha_{si}L\Pi_0+m_i D}{2\alpha_{si}\Pi_0 L+D+\eta_i L}$, where $i = 1, 2$. Note that the oncotic pressure difference is zero across the epithelial cells ($\Delta\Pi_p = 0$) while it is non-zero across the endothelial cells ($\Delta\Pi_p = \Pi_p$).

In Fig. 5 and Fig. 7-10, when not specified, the energy inputs for kidney epithelial pump and endothelial pump are: $(\tilde{G}_{a1}, \tilde{G}_{a2}) = (29.10, 11.64)$. All other parameters are the same for both pumps (See Table 1,2). For the filter calculation in Fig. 4-5, we assume that the flux only depends on the pressure difference and the flux curve is solved with zero osmolarity difference and zero energy input ($\Delta\Pi = \Delta\Pi_P = 0, \tilde{G}_a = 0$). Notably, we have assumed that the endothelial tissue is actively pumping the same as the epithelial tissue, which has not been established convincingly in experiments. The degree of "active pumping" can be adjusted by tuning the active transport coefficient $\gamma_2$ or $\tilde{G}_{a2}$. Results when $\tilde{G}_{a2} = 0$ are shown in Fig. 10.

#### Supplementary Note 3: Parameter estimation

For an isolated pump, the most important parameters are related to passive and active ion transport. In kidneys, a cell in the distal convoluted tubule has up to 50 million pumps [8]. An important pump for sodium transport is the ATP hydrolysis-driven sodium-potassium pump, which exchanges three intracellular $Na^+$ ions for two extracellular $K^+$ ions 100 times per second [9]. The maximum ion pumping rate can then be estimated as: $J_{total} = 3 \times 100 \times 5 \times 10^7 = 1.5 \times 10^{10}$ s$^{-1}$. Assume that the cell is cylindrical and the radius is 10 $\mu$m, the sodium flux can be estimated as: $J = \frac{1.5\times10^{10}/6.02\times10^{23}}{4\pi(10^{-5})^2} \approx 2 \times 10^{-5}$ mol/(m$^2\cdot$ s). This estimation provides an upper limit for the pumping rate. In general, the pumping rate is in the range of $J = 10^{-7} \sim 10^{-5}$ mol/(m$^2\cdot$ s) [1]. We assume that the passive ion transport rate is the same as that of active pumping and the osmotic pressure difference across cell membrane is $\Delta\Pi \approx 100$ Pa. The coefficient of passive ion transport through ion channels can be calculated as $\beta = J/\Delta\Pi \times RT \approx 2.8 \times 10^{-11} \sim 2.8 \times 10^{-9}$ m/s. Since we directly deal with osmotic pressure instead of concentration, we have a modifying term $RT$ in the estimation that converts the concentration into osmotic pressure. The coefficient of active ion pumping is estimated as follows. First, the energy input for ion pumping typically comes from ATP hydrolysis, which takes the value around 30 kJ/mol and the dimensionless energy input is $\tilde{G} = G/RT \approx 11.64$. The active pumping rate can be expressed as: $J_a = \gamma\tilde{G}(\Pi_{m1} - \frac{\Delta\Pi_m}{e^{\tilde{G}}-1}) \approx \gamma\tilde{G}\Pi_{m1}$, The coefficient is then calculated as: $\gamma = J/(\tilde{G}\Pi_{m1}) \times RT \approx 2.6 \times 10^{-6} \sim 2.6 \times 10^{-4}$ m/s. We can also estimate the coefficients of solute flux dependence on pressure ($m$) and osmolarity ($m'$). Theoretically, the total energy input should be zero under stall pressure (We assume it is around 500 Pa). Therefore, $m \approx \gamma\tilde{G}_a\Pi/500 = 5.2 \times 10^{-5} J/(Pa \cdot s)$. Similarly, $m' \approx 5.2 \times 10^{-5} J/(Pa \cdot s)$.

For the whole circulatory system, the important parameters are resistances of different parts. Since we focus on the static property, we will neglect the capacitors. In our model, the system is modeled as a simplified version of paper [7]. All the resistances are estimated based on this work. Differently, the resistances of aorta and vena cava in our model are simplified to be concentrated instead of distributed. The resistance of the glomerulus is estimated as follows. The glomerular capillary pressure is about 55 mmHg and hydrostatic pressure within Bowman's space is 20 mmHg. The osmotic pressure difference across glomerulus is 28 mmHg [10]. The kidneys filter 180 L fluid per day [11]. Therefore, the glomerulus resistance is $R_G = (\Delta P - \Delta \Pi)/I = \frac{(55-20)-28}{180/24/3600} = 3.36\ \mathrm{mmHg} \cdot \mathrm{s} \cdot \mathrm{mL}^{-1}$.

### Supplementary Tables

Table 1: Glossary of model variables and their numerical estimates for the pumping element. See text for references.

| Symbol | Description | Values | Source |
|---|---|---|---|
| $D$ | Diffusion coefficient of ions, e.g. $K^+$ ($m^2$/s) | $2 \times 10^{-9}$ | [3, 4] |
| $\gamma$ | Active ion transport coefficient (m/s) | $2.8 \times 10^{-9}$ | [1, 8, 9] |
| $\beta$ | Passive ion transport coefficient (m/s) | $2.6 \times 10^{-4}$ | [1, 8, 9] |
| $\alpha$ | rate constant of water transport ($m \cdot s^{-1} \cdot Pa^{-1}$) | $10^{-9}$ | [1] |
| $G$ | Energy input of active ion pumping (kJ/mol) | 30 | [1] |
| $L$ | Thickness of renal epithelial (endothelial) tissue (m) | $10^{-5}$ | - |
| $m$ | proportionality constant between ion flux and pressure gradient (J/ (Pa·s)) | $5.2 \times 10^{-5}$ | - |
| $m'$ | proportionality constant between ion flux and osmolarity gradient (J/ (Pa·s)) | $5.2 \times 10^{-5}$ | - |
| $S_{Ep}$ | Surface area of the renal tubule ($m^2$) | 13.276 | [12, 13, 14, 2] |
| $S_{Ed}$ | Surface area of the endothelial tissue in renal veins ($m^2$) | 0.6 | [15] |

Table 2: Glossary of model variables and their numerical estimates for the circulatory system. See text for references.

| Symbol | Description | Values | Source |
|---|---|---|---|
| $R_A$ | Effective resistance of the aorta (Pa·s/$m^3$) | $7.78 \times 10^7$ | [7] |
| $R_V$ | Effective resistance of the inferior vena cava (Pa·s/$m^3$) | $9.98 \times 10^6$ | [7] |
| $R_O$ | Effective resistance of the capillaries in all organs (Pa·s/$m^3$) | $1.34 \times 10^8$ | [7] |
| $R_G$ | Effective resistance of the glomerulus (Pa·s/$m^3$) | $4.47 \times 10^8$ | [7] |
| $P_s$ | Pressure generated by the heart (Pa) | $1.6 \times 10^4$ | [7] |
| $\Pi_1$ | Total blood osmolarity (Pa) | $8 \times 10^5$ | [16] |
| $\Pi_p$ | Blood oncotic pressure (Pa) | $3.8 \times 10^3$ | [10] |

Table 3: Expressions of some effective model parameters. See text for references.

| Symbol | Description | Expression |
|---|---|---|
| $r$ | dimensionless energy input | $r = \frac{\gamma}{\eta}\frac{\Delta G_a}{RT}$ |
| $\theta$ | Model coefficient | $\theta = \frac{r}{r+2}$ |
| $\Pi_0$ | Weighted mean osmolarity around the pumping element | $\Pi_0 = \frac{(r+1)\Pi_a + \Pi_b}{r+2}$ |
| $\Pi_{02}$ | Effective osmolarity parameter | $\Pi_{02} = \frac{(r+1)^2\Pi_a - \Pi_b}{r+2}$ |
| $\alpha_s$ | First effective water permeation constant | $\alpha_s = \frac{\alpha}{8\mu L\alpha/R_{cell}^2 + 2}$ |
| $\alpha_{ss}$ | Second effective water permeation constant | $\alpha_{ss} = \frac{\alpha_s(D+\eta L)}{2\alpha_s\Pi_0 + (D+\eta L)}$ |

### Supplementary Figures

#### Cell monolayer fluid and solute pumping model

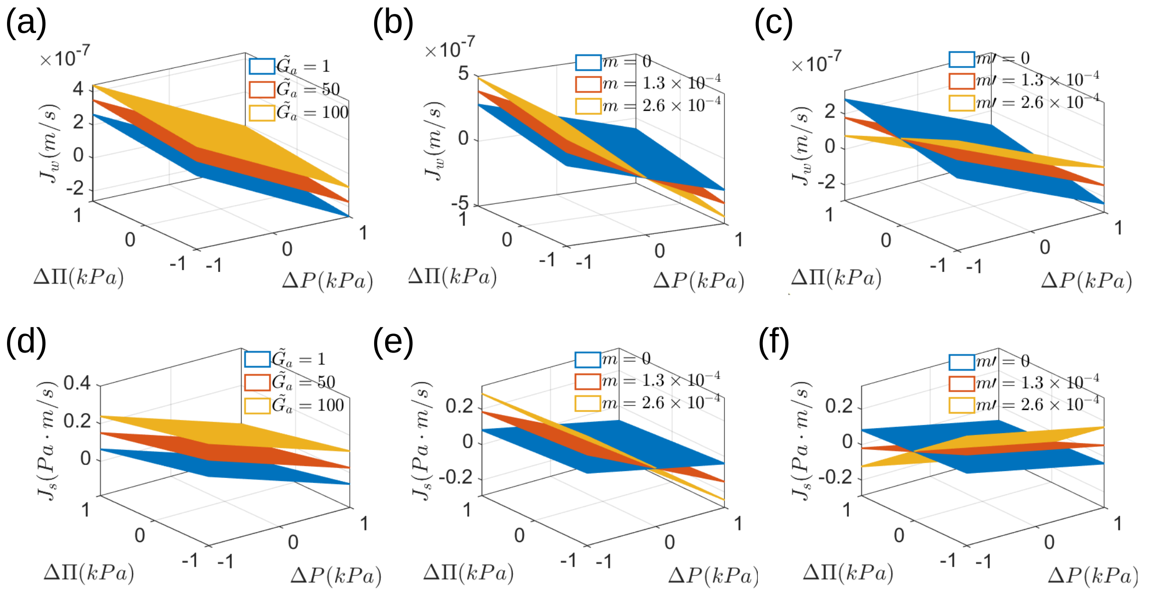


Figure 2: Generalized "pump performance surface" for an isolated pump: Dependence of water (solute) flux on basal-apical pressure difference ($\Delta P$) and osmolarity difference ($\Delta\Pi$) with different energy input ($\tilde{G}_a$) and solute flux dependence on pressure and osmolarity ($m, m'$). The energy input $\tilde{G}_a$ increases both the water and solute flux (a&d). In the generalized pump performance surface, $m, m'$ decrease the slope of the fluxes with respect to pressure gradient $\Delta P$ and osmolarity gradient $\Delta\Pi$ (b,c,e,f).

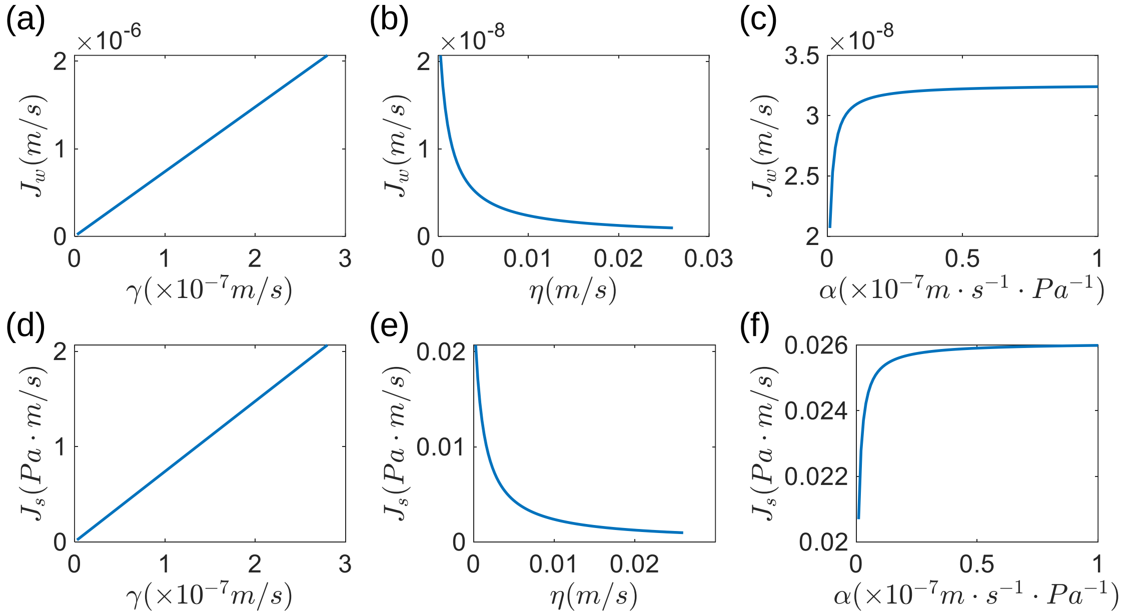


Figure 3: Influence of active, passive ion transport coefficients $(\gamma, \eta)$ and water permeability of the membrane $(\alpha)$ on water (a-c) and solute flux (d-f) for an isolated pump. (a) Active transport of ion increases the water flux. (b) Increasing passive transport coefficient decreases water flux. (c) Increase in water permeability leads to increased water flux, reaching a plateau. (d-f) Results on solute flux.

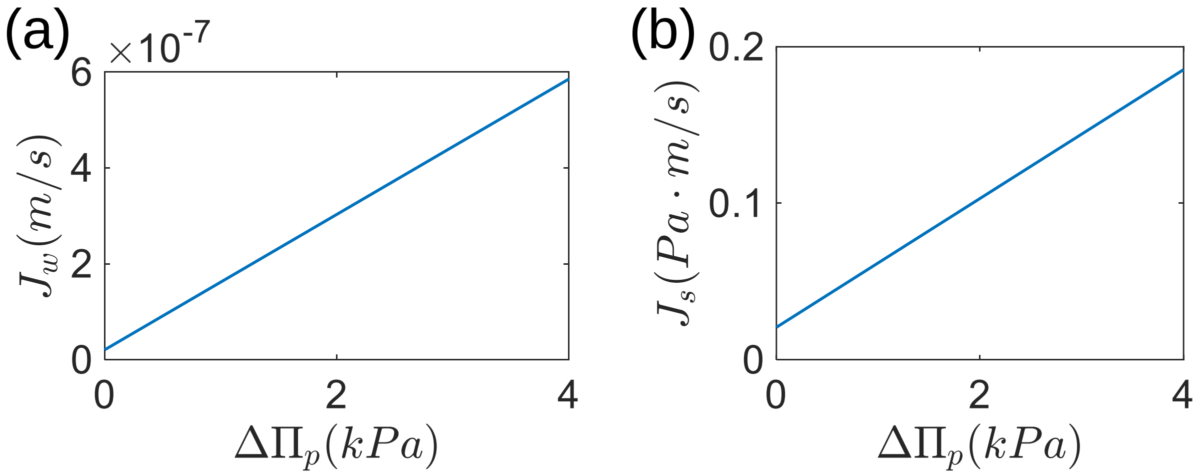


Figure 4: Influence of macromolecule osmotic pressure gradient $\Delta\Pi_p$ on water (a) and solute flux (b) for an isolated pump. Both water and solute fluxes increase with the osmotic pressure gradient.

### A one-pump circulation model

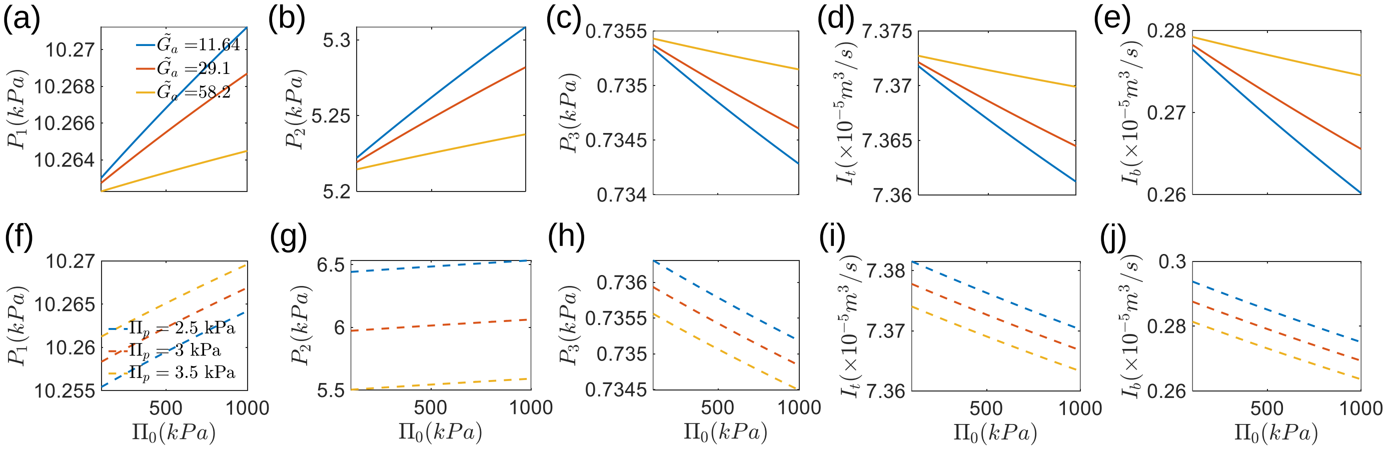


Figure 5: Influence of osmotic pressure on node pressure and blood flux with different energy input for ion pump ($\tilde{G}_a$,a-e) and blood plasma oncotic pressure ($\Pi_p$,f-j). (a)-(c) Increase of the external osmotic pressure causes decrease in pressure at node 3 and increase in node 1 and 2. Increase of energy input decreases the pressure at node 1 and 2 while increases the pressure at node 3. (d)-(e) Total blood flux and the branch flux across the pump both decrease with external osmolarity while increase with energy input. (f)-(h) Oncotic pressure in blood plasma increases the pressure at node 1 while decreases the pressure at node 2 and 3. (i)-(j) Total blood flux and the branch flux across the pump both decrease with the increase of blood oncotic pressure. In (f)-(j), the energy input is set as: $\tilde{G}_a = 11.64$. The water transport constant is set as $\alpha = 5 \times 10^{-11}\ m \cdot s^{-1} \cdot Pa^{-1}$.

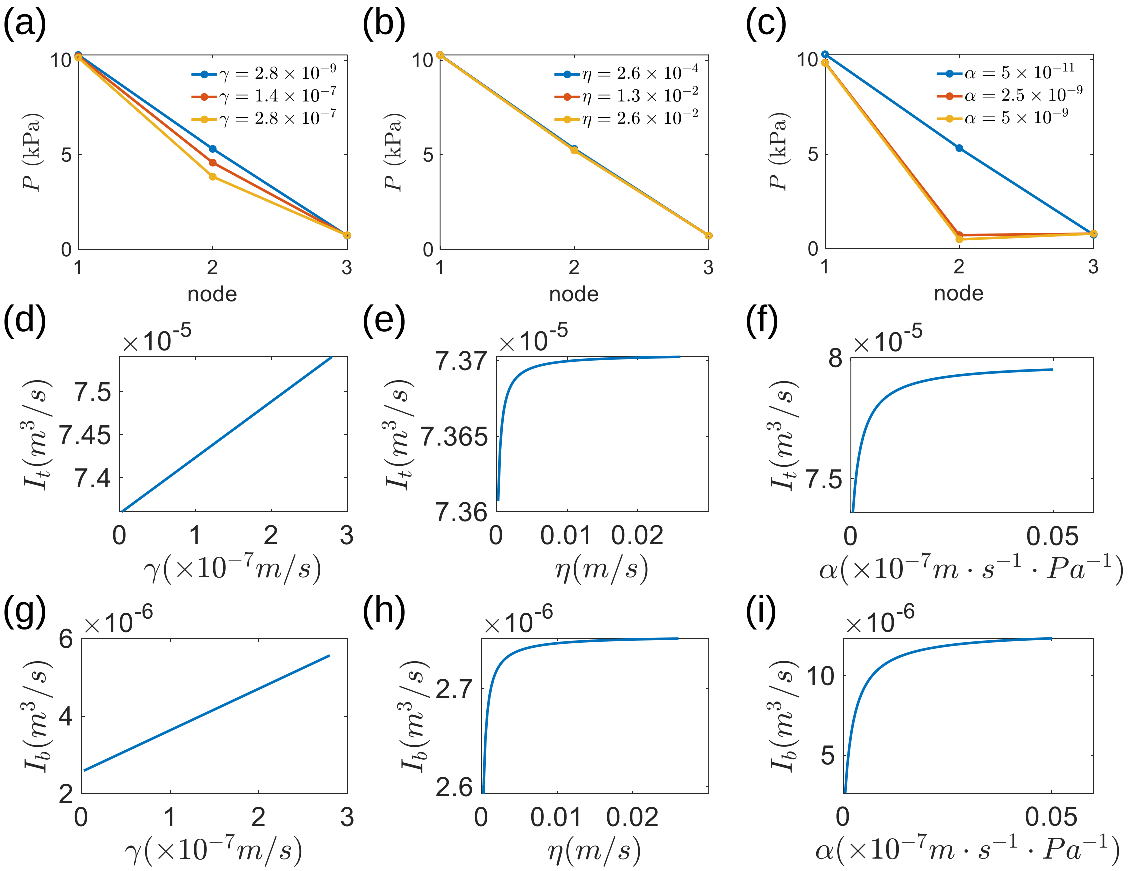


Figure 6: Influence of active, passive ion transport coefficients $(\gamma, \eta)$ and water permeability of the membrane $(\alpha)$ on pressure distribution (a-c), total blood flux (d-f) and branch flux (g-i) across the pumping element for the one-pump network. (a)-(c) Both the active ion transport coefficient $(\gamma)$ and water permeability $(\alpha)$ of the pump decrease the pressure in the lumen of the renal tubule (node 2). (d)-(i) Both the total flux and branch flux across the pump increase with $\gamma, \eta$, and $\alpha$.

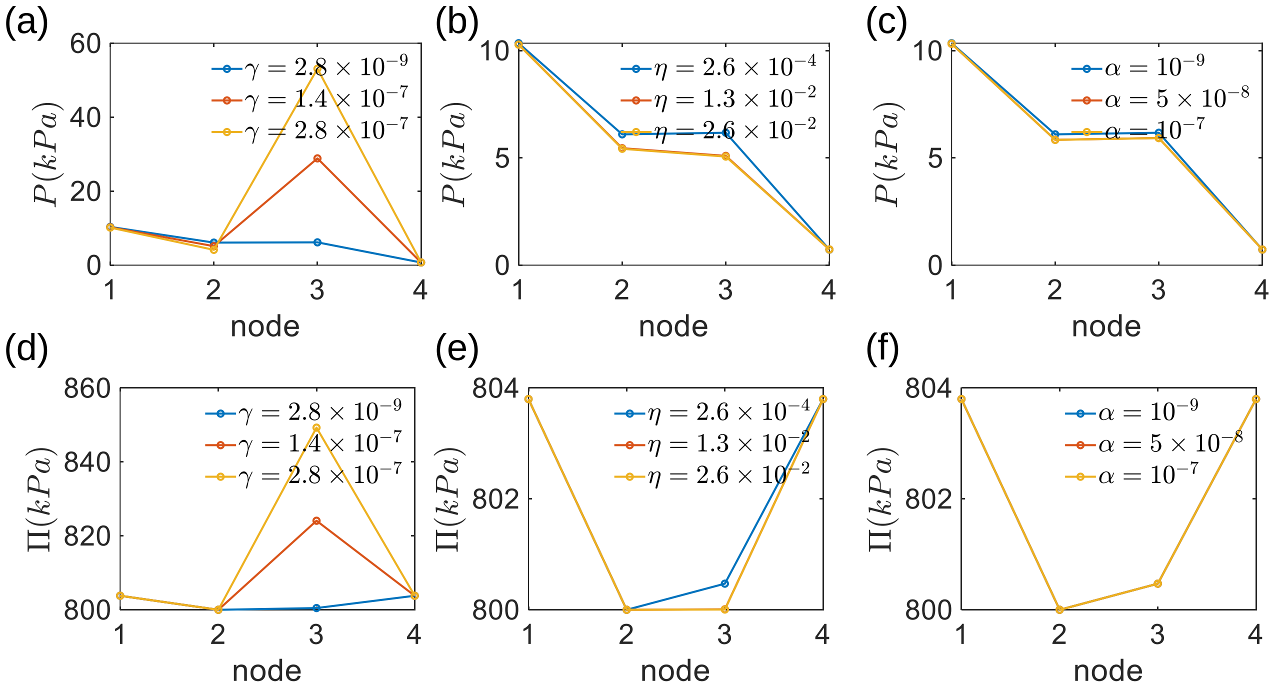


Figure 7: (a)-(c) Influence of active, passive ion transport coefficients $(\gamma, \eta)$ and water permeability of the membrane $(\alpha)$ on pressure distribution. The interstitial pressure (node 3) increases with the increase of active ion transport coefficient $\gamma$ and decrease of passive transport coefficient $\eta$ and water permeability $\alpha$. (d)-(f) Results on osmolarity distribution for the two-pump network. The interstitial osmolarity (node 3) increases with the increase of active ion transport coefficient $\gamma$ and decrease of passive transport coefficient $\eta$.

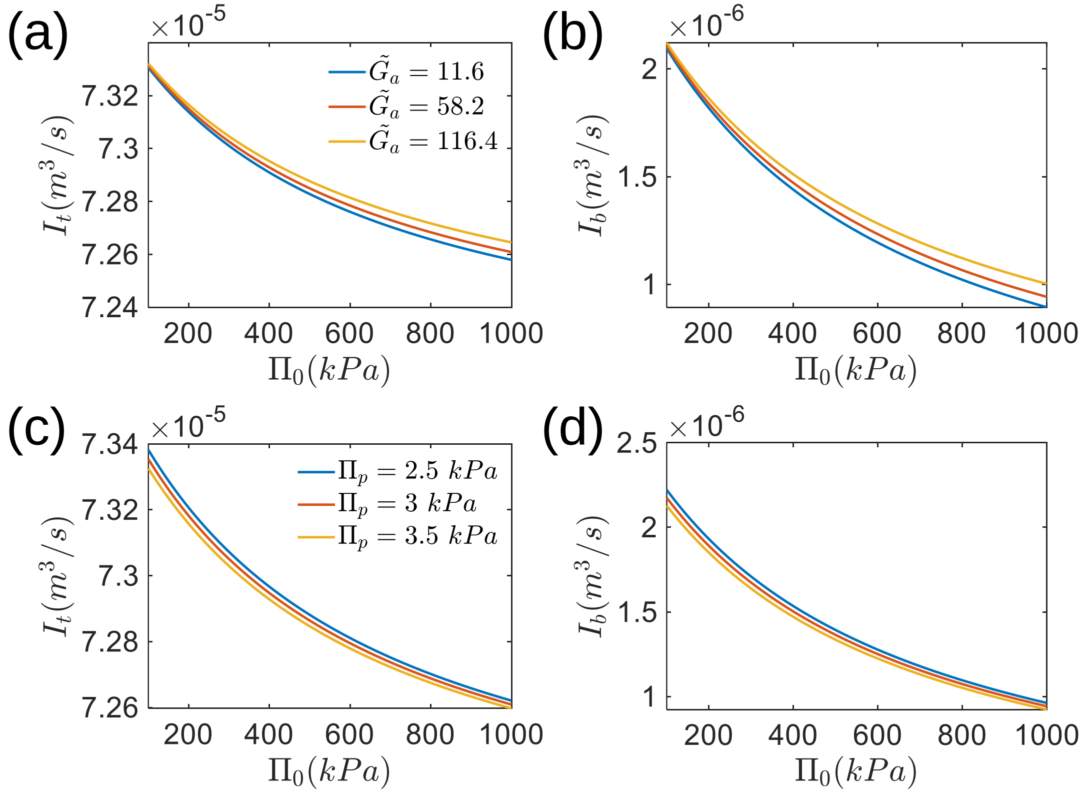


Figure 8: Influence of osmotic pressure on total flux and branch flux across the pumps with different ion pump energy input and blood oncotic pressure. According to the two-pump model, both the total flux and branch flux across the pumping element decrease with elevated external osmotic pressure. (a)-(b) Increase of energy input for the ion pump increases the blood flux. (c)-(d) Increase of blood plasma oncotic pressure decreases the blood flux.

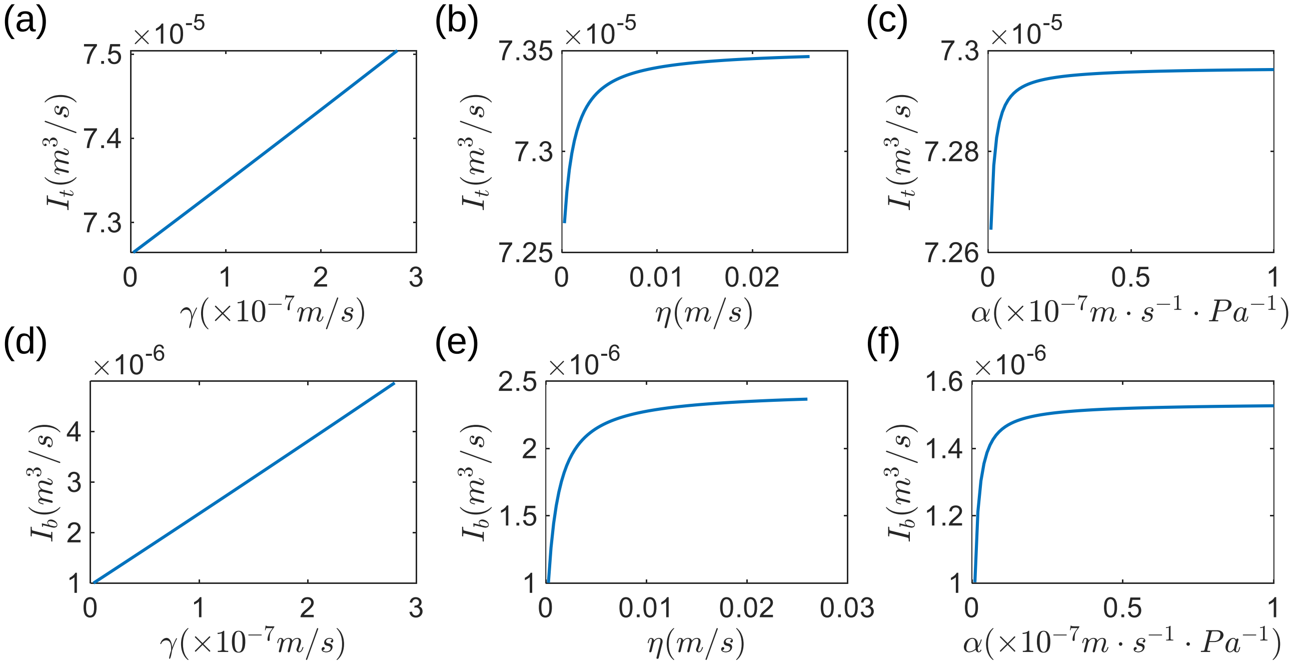


Figure 9: Influence of active, passive ion transport coefficients $(\gamma, \eta)$ and water permeability of the membrane $(\alpha)$ on total blood flux (a-c) and branch flux (d-f) across the pumping elements for the two-pump network.

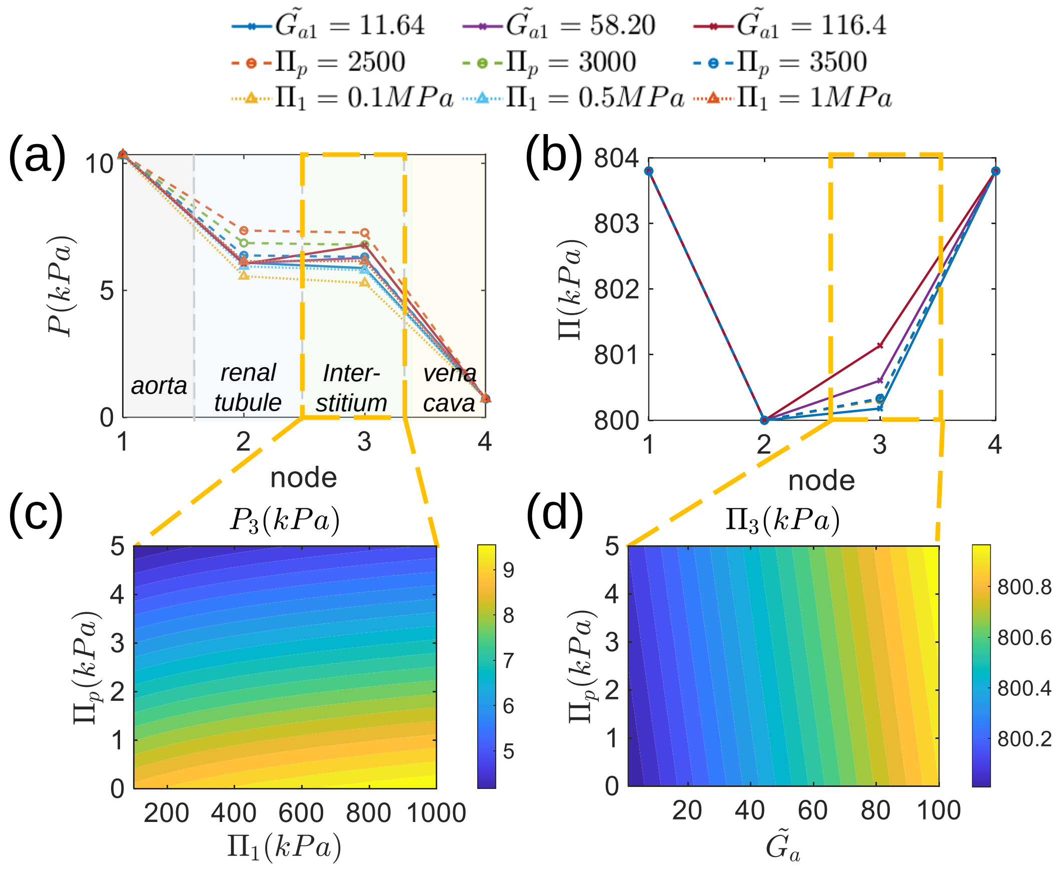


Figure 10: Influence of total blood osmotic pressure, oncotic pressure and energy input on hydraulic pressure and osmotic pressure when endothelial cells are not actively pumping ($\tilde{G_{a2}} = 0$). (a)-(b) spatial distribution of pressure and osmolarity with different energy input, blood oncotic pressure and total osmolarity in blood plasma. (c)-(d) Influence of total blood osmotic pressure, oncotic pressure and energy input on hydraulic pressure and osmotic pressure in the interstitium. When not specified, the energy inputs for kidney epithelial pump and endothelial pump are: $(\tilde{G_{a1}}, \tilde{G_{a2}}) = (29.10, 0)$. All other parameters are the same for both pumps.

### References

[1] Jiang H, Sun SX. Cellular pressure and volume regulation and implications for cell mechanics. *Biophys. J.* 2013; 105(3):609-19.

[2] Layton AT, Layton HE. A computational model of epithelial solute and water transport along a human nephron. *PLoS Comput. Biol.* 2019; 15(2):e1006108.

[3] Samson E, Marchand J, Snyder KA. Calculation of ionic diffusion coefficients on the basis of migration test results. *Mater Struct* 2003: 156-65.

[4] Teng X, Huang Q, Dharmawardhana CC, Ichiye T. Diffusion of aqueous solutions of ionic, zwitterionic, and polar solutes. *J. Chem. Phys.* 2018; 148(22).

[5] Choudhury MI, Li Y, Mistriotis P, Vasconcelos AC, Dixon EE, Yang J, Benson M, Maity D, Walker R, Martin L, Koroma F. Kidney epithelial cells are active mechano-biological fluid pumps. *Nat. Commun.* 2022; 13(1):2317.

[6] Parsons DS, Paterson CR. Fluid and solute transport across rat colonic mucosa. *Q. J. Exp. Physiol. Cogn. Med. Sci.* 1965; 50(2):220-31.

[7] Kung E, Pennati G, Migliavacca F, Hsia TY, Figliola R, Marsden A, Giardini A, MOCHA Investigators. A simulation protocol for exercise physiology in Fontan patients using a closed loop lumped-parameter model. *J. Biomech. Eng.* 2014; 136(8):081007.

[8] Clausen MV, Hilbers F, Poulsen H. The structure and function of the Na, K-ATPase isoforms in health and disease. *Front. physiol.* 2017 Jun 6;8:371.

[9] Gadsby DC. Spot the difference. *Nature.* 2004; 427(6977):795-7.

[10] Feher JJ. Quantitative human physiology: an introduction. *Academic press.* 2017 Jan 2.

[11] Kaufman DP, Basit H, Knohl SJ. Physiology, glomerular filtration rate. 2018.

[12] Chevalier RL. The proximal tubule is the primary target of injury and progression of kidney disease: role of the glomerulotubular junction. *Am. J. Physiol. Renal Physiol.* 2016;311(1):F145-61.

[13] Aličelebić S. Proximal convoluted tubules of the rats kidney–a stereological analysis. *Biomol. Biomed.* 2003; 3(1):36-9.

[14] Bertram JF, Douglas-Denton RN, Diouf B, Hughson MD, Hoy WE. Human nephron number: implications for health and disease. *Pediatr. Nephrol.* 2011; 26:1529-33.

[15] Bohle A, Aeikens B, Eenboom A, Fronholt L, Plate WR, Xiao JC, Greschniok A, Wehrmann M. Human glomerular structure under normal conditions and in isolated glomerular disease. *Kidney Int.* 1998; 54:S186-8.

[16] Levenbrown Y, Costarino AT. Edema. Nephrology and Fluid/Electrolyte Physiology. 2019; Elsevier.
